# FOXA2 potently represses viral gene expression by targeting NF-κB–dependent transcription in liver cells

**DOI:** 10.64898/2026.01.27.701993

**Authors:** Tariq Suleiman, Michael D. Walker

**Affiliations:** Department of Biomolecular Sciences, Weizmann Institute of Science, Rehovot 76100, Israel

**Keywords:** transcription factor, transcription repressor, promoter, viral transcription, NF-κB, liver, gene regulation

## Abstract

The transcription factor Forkhead box A2 (FOXA2) is a master regulator of endoderm development and of mature endoderm-derived organs, including the liver, and tight control of its expression and activity is paramount for the proper execution of developmental and homeostatic gene programs. Unexpectedly, despite its established role as a pioneer transcription factor and transcriptional activator, FOXA2 exhibited robust repressive activity in differentiated liver cells. FOXA2 over-expression led to widespread repression of endogenous genes, including autorepression of the FOXA2 gene itself, and, notably, to dramatic suppression of viral regulatory elements. Among these, the SV40 early promoter—an essential driver of viral replication and cellular transformation—was strongly repressed by FOXA2. DNA binding by FOXA2 is sufficient to drive repression of the SV40 early promoter. Repression was most pronounced on regulatory elements whose activity is highly dependent on NF-κB, identifying NF-κB–driven promoters as a key target of FOXA2 repressive activity.

Mechanistically, EMSA and immunoblot analyses indicate that FOXA2 does not repress viral promoters through direct competition with NF-κB for DNA binding. Instead, FOXA2 expression was associated with reduced NF-κB protein abundance, at least in part through proteasome-dependent degradation, revealing a novel indirect mechanism by which FOXA2 constrains NF-κB–dependent transcription. Given that many viruses, including SV40 and HIV-1, rely heavily on NF-κB activity to drive early gene expression and the transition from latency to productive infection, our findings suggest that FOXA2 may function as a host-derived, context-dependent restriction factor that limits viral gene expression in differentiated tissues.

## INTRODUCTION

Forkhead box A2 (FOXA2) protein, formerly known as hepatocellular nuclear factor 3-beta (HNF3β), belongs to the Forkhead box A (FOXA) subfamily of transcription factors (TFs), which also includes FOXA1 (HNF3α) and FOXA3 (HNF3γ) in mammals. Members of this family contain a conserved DNA-binding domain (DBD), the Forkhead or winged-helix domain, which mediates sequence-specific DNA recognition [1]. In addition, FOXA2 contains transactivation domains (TADs) at both its N- and C-termini, enabling interactions with a variety of transcriptional coregulators [2]. FOXA2 is the earliest expressed member of the FOXA family during embryogenesis and plays a central role in coordinating precisely timed signaling events and hierarchical gene expression programs required for endoderm specification [3]. Consistent with this role, FOXA2 knockdown was shown to drive genome-wide changes and impair cellular differentiation [4], and genetic ablation of FOXA2 in mice results in severe defects in endoderm formation [5]. Beyond development, FOXA2 remains essential for the function and maintenance of mature endoderm-derived organs, including the liver [6]. Together, these findings establish FOXA2 as a master regulator of gene expression, underscoring the importance of tight control over its expression and activity.

Mechanistically, FOXA2 functions as a pioneer transcription factor that can bind compacted chromatin and interact with core histones, thereby facilitating chromatin opening and enabling the recruitment of additional transcription factors [3,7,8]. Through this activity, FOXA2 regulates diverse transcriptional programs across tissues. In the pancreas, FOXA2 activates expression of PDX1, a key regulator gene of β-cell identity, thereby permitting subsequent activation of the insulin gene [9]. In the liver, FOXA2 controls metabolic gene networks involved in glucose and lipid homeostasis, including genes encoding the glucocorticoid receptor (GR), glucose transporter 2 (GLUT2), and glucose-6-phosphatase (G6P), among others [3,10,11].

FOXA2 has been predominantly characterized as a transcriptional activator. However, accumulating evidence indicates that FOXA2 can also function as a transcriptional repressor in specific contexts. For example, FOXA2 negatively regulates expression of the ABCA1 gene in hepatocytes [12], and over-expression of FOXA2 represses transcription of the hepatic transporter gene Slco1b3 [13]. Additionally, it has been reported that in the adult liver, FOXA2’s function is not restricted to maintaining enhancer activity and opening the chromatin during early development [14,15]. Together, these observations suggest that the regulatory repertoire of FOXA2 extends beyond transcriptional activation, although the biological scope and mechanistic basis of its repressive activity remain poorly understood.

Despite extensive characterization of FOXA2 target genes, comparatively little is known about the mechanisms governing transcriptional regulation of the FOXA2 gene itself. A limited number of factors have been implicated, including repression by Tcf7l1 and microRNA-124a [16,17]. In addition, FOXA2 protein activity and stability are modulated by post-translational modifications such as phosphorylation by AKT/PKB and IKKα [18,19]. However, how transcriptional initiation at the FOXA2 locus is regulated, and whether alternative regulatory elements contribute to its expression, remains largely unexplored. One potential mechanism contributing to this regulatory complexity is the use of alternative transcription start sites. Indeed, inspection of the FOXA2 genomic locus, using the UCSC Genome Browser, revealed the presence of two alternative transcripts (Fig. 1A). The canonical transcript, which has been described in the literature, initiates from an upstream promoter and is composed of three exons. In contrast, a shorter and incompletely characterized transcript initiates from a downstream genomic region and consists of two exons. Hereafter, the upstream canonical promoter and its associated transcript are referred to as P1, whereas the downstream transcript and its putative promoter are referred to as P2. The predicted protein isoform encoded by the P2 transcript contains an N-terminal 6 amino acid extension absent from the P1-encoded isoform (Fig. 1B). Distinct first exons suggests transcription driven from alternative promoters [20]. We validated the function of the putative promoter using reporter gene experiments (Fig 1C). The use of alternative promoters is a common mechanism for achieving regulatory complexity in mammalian genomes, with approximately half of human protein-coding genes employing alternative transcription start sites [20]. This strategy is particularly prevalent among genes involved in transcriptional regulation and development, where alternative promoters can confer distinct temporal or tissue-specific expression patterns or generate protein isoforms with divergent functional properties [21,22]. These observations prompted us to investigate whether alternative promoter usage at the FOXA2 locus produces isoforms with distinct functional activities.

**Figure 1.**
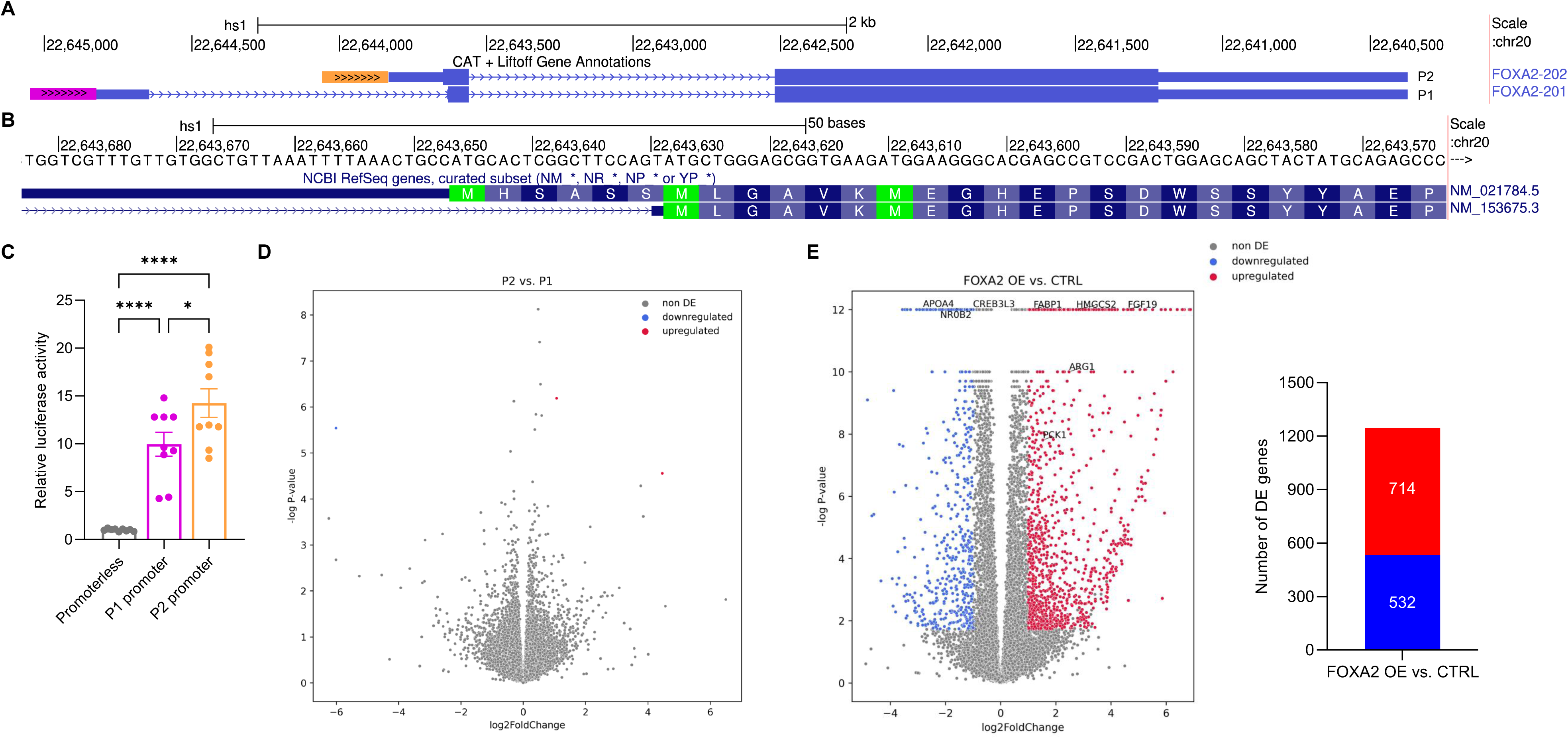
The FOXA2 locus generates two closely similar protein isoforms in HepG2 cells. **(A)** Schematic representation of the human FOXA2 locus based on the UCSC Genome Browser. The FOXA2 locus encodes two alternative transcripts with distinct transcription start sites: the canonical transcript (P1, lower), containing three exons (blue bars), the first of which is noncoding, and an alternative transcript (P2, upper) containing two coding exons. Thick bars indicate coding sequences, whereas thin bars indicate noncoding regions. The location of the canonical promoter (FOXA2-P1) is shown in magenta, and the alternative promoter (FOXA2-P2) is shown in orange. **(B)** DNA and amino acid sequences corresponding to the N termini of the protein isoforms encoded by P1 (lower) and P2 (upper) FOXA2 transcripts. NCBI RefSeq transcript accession numbers corresponding to each isoform are indicated to the right. **(C)** Activity of the FOXA2 promoters. HepG2 cells were transiently transfected with 450 ng of firefly luciferase reporter plasmids containing either the P1 or P2 promoter, or with a promoterless firefly luciferase vector (pGL3-Basic) as a control. Results are presented as mean ± SEM from three independent experiments, each with 2–4 biological replicates. Statistical analysis was performed using ordinary one-way ANOVA with Brown–Forsythe correction (****p < 0.0001; *p < 0.05). **(D, E)** RNA-seq analysis of HepG2 cells transfected with FOXA2 isoforms. Five biological replicates were transfected with empty vector (EV; control), FOXA2 P1 expression plasmid, FOXA2 P2 expression plasmid, or both P1 and P2 expression plasmids (each comprising half of the total DNA amount). Volcano plots depict differential gene expression, with P1-transfected samples used as the reference in (D) and control samples used as the reference in (E, left). Eight selected differentially expressed genes are highlighted in (E, left). The number of differentially expressed (DE) genes in comparisons between FOXA2- or EV-transfected cells is indicated in (E, right). Threshold for significant differential expression was set at adjusted p ≤ 0.05 and |log₂ fold change| ≥ 1.

Our experiments indicated that the isoforms have equivalent biological function. Unexpectedly, however, FOXA2 expression in liver cells led to transcriptional repression of multiple cellular genes and particularly robust repression of viral gene promoters. Repression was most pronounced on regulatory elements whose activity is highly dependent on NF-κB, e.g. the simian virus 40 (SV40) early promoter, and appears to function indirectly by reducing NF-κB protein abundance.

## RESULTS

### FOXA2 isoforms exhibit similar activity and mediate broad gene repression

We used reporter gene experiments to test directly the promoter activity of the putative P2 promoter. Indeed, we observed that this genomic region displayed significant activity that was even higher than that of the canonical P1 promoter (Fig 1C). Consistent with this, using RT-PCR analysis with transcript-specific primers, we were able to confirm that both promoters drive endogenous expression of their respective transcripts and that the putative P2 promoter has similar or even higher activity to that of P1 in HepG2 liver cells (data not shown). To identify possible functional differences between the two protein isoforms, we initially sought to perform loss-of-function experiments by selectively downregulating each variant, either by siRNA-mediated knockdown or by CRISPR-Cas9-mediated deletion of the individual promoters. However, both approaches proved technically challenging and impractical. We therefore conducted an over-expression experiment by introducing expression plasmids encoding each FOXA2 isoform in HepG2 cells. To allow isolation of only successfully transfected cells, cells were co-transfected with a GFP-encoding plasmid at a concentration we determined empirically to be minimally toxic. We isolated RNA from GFP-positive FACS-sorted cells and performed RNA-seq analysis using the MARS-seq protocol [23,24]. Over-expression of P1 and P2 isoforms elicited highly similar transcriptional responses (Fig.1D). The small number of differentially expressed (DE) genes observed between the two conditions could be attributed solely to outliers within the biological replicates (i.e. one sample out of five), indicating no isoform-specific effects. Thus, we found no evidence for functional differences between the two FOXA2 isoforms. On the other hand, expression of either FOXA2 isoform markedly altered global gene expression (Fig. 1E), with a bias toward transcriptional activation, including several previously reported target genes highlighted in the volcano plot (Fig. 1E, red) [25–27]. Nevertheless, a substantial number of genes were also repressed (Fig. 1E, blue), with over 500 downregulated compared to ∼700 upregulated genes. This finding contrasts with the prevailing view of FOXA2 as primarily a positive regulator of gene expression [3]. A list of the most strongly repressed genes, along with associated enriched biological processes, is provided in Table S1 & Fig. S1 respectively.

### Repression of endogenous genes is corroborated by transactivation assays

In order to validate this unexpectedly broad repression, we conducted transactivation assays using reporter plasmids harboring several gene promoters upstream of a firefly luciferase reporter (Fig. 2A). Four genes, including FOXA2 itself, that showed statistically significant repression upon FOXA2 over-expression in the RNA-seq analysis (Fig. 2B) were selected. None of these are established FOXA2 targets, although p53 upregulated modulator of apoptosis (PUMA) has been reported to be bound by FOXA2 in vivo based on publicly available ChIP-seq datasets (NCBI Sequence Read Archive accessions SRX100448, SRX10475773, SRX2368886, SRX3322258/9). The transcriptional activity of these promoters was assessed in the presence or absence of over-expression of the FOXA2 protein isoforms. Overall, the transactivation assays closely mirrored the RNA-seq results, with similar trends and magnitudes of repression (Fig. 2C). For example, the pro-apoptotic gene PUMA that exhibited an approximately 40% reduction in endogenous expression, displayed ∼70% decrease in activity of the exogenous promoter in the reporter assay. Similarly, in both experimental systems, TP53, a positive regulator of PUMA [28], showed 40-50% repression upon FOXA2 over-expression, while CDK4 displayed an approximately 30% reduction. Collectively, these findings validate the RNA-seq data and point to a potential role for FOXA2 in repressing the transcription of genes relevant to tumorigenesis.

**Figure 2.**
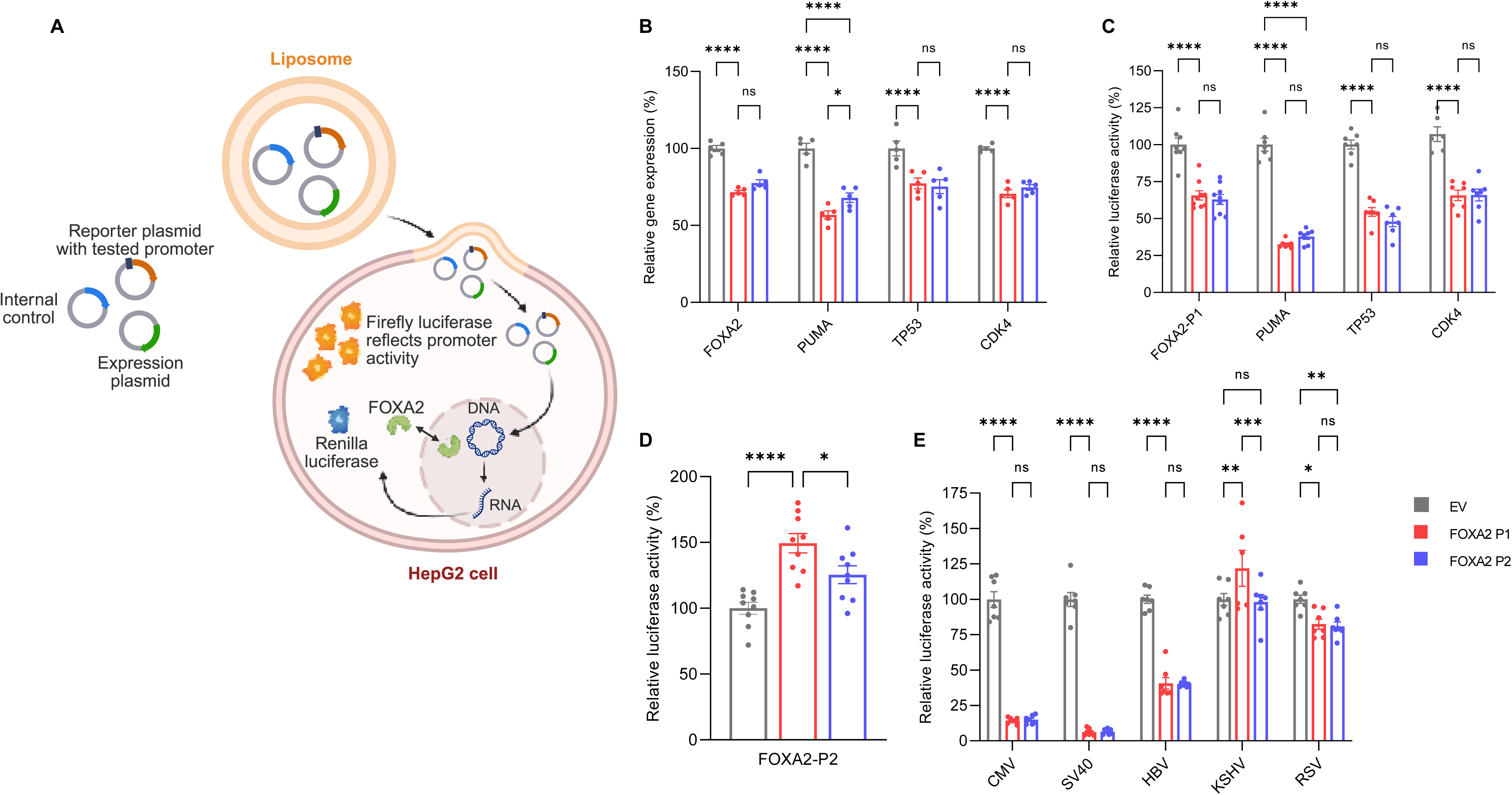
FOXA2 represses cellular and viral regulatory elements in HepG2 cells, including its own expression. **(A)** Schematic representation of the luciferase-based transactivation system used to assess regulatory element activity under FOXA2 over-expression conditions (created with BioRender): a reporter plasmid harboring the tested regulatory elements upstream of the firefly luciferase gene (orange), a plasmid encoding Renilla luciferase used as an internal control (blue), and an expression plasmid encoding a transcription factor (green; FOXA2 in this case). **(B)** Relative expression levels of selected endogenous genes derived from RNA-seq analysis, shown with comparisons among the indicated groups. Results are presented as mean ± SEM. **(C–E)** Transactivation assays of regulatory elements by FOXA2. HepG2 cells were transiently transfected with 200 ng of plasmids containing the indicated cellular (C, D) or viral (E) regulatory elements upstream of the firefly luciferase reporter gene. In addition, 90–350 ng of either noncoding empty vector (EV; control), plasmid encoding FOXA2 P1, or plasmid encoding FOXA2 P2 were co-transfected. The same total amount of DNA was used in all transfections (with EV used as carrier to reach a total of 600 ng when required). Results are presented as mean ± SEM for three independent experiments with 2–3 biological replicates per experiment. Statistical analysis was performed using ordinary two-way ANOVA with Tukey’s multiple comparisons test (B, C, E) or one-way ANOVA with Brown–Forsythe correction (D). (****p < 0.0001, ***p < 0.001, **p < 0.01, *p < 0.05, ns p > 0.05).

### FOXA2 exhibits auto-repression in liver cells

Auto-regulation is common in genes encoding TFs, and serves as an important mechanism to refine and maintain accurate TF levels in accordance with cellular demands at any given time. Master regulator genes are recognized for their role in positive feedback loops, particularly in development where they are capable of upregulating their own expression to drive irreversible cell fate decisions [29]. In light of this paradigm, the observed repression of FOXA2 activity at its canonical P1 promoter was unexpected. Transcription factors, including FOXA2, are known for their sensitivity to copy number changes and their tendency to function in a dose-dependent manner [30]. To exclude the possibility that FOXA2 regulates the tested gene promoters in a bi-phasic, dose-dependent manner, all transactivation assays were performed using a range of FOXA2 expression levels. Across all promoters examined, the same regulatory pattern was observed, with repression evident at both low and high FOXA2 protein amounts (data not shown). These findings argue against biphasic regulation and instead indicate that repression of FOXA2’s canonical promoter is not dependent on FOXA2 dosage. We next considered whether this regulatory behavior could reflect differential control of alternative FOXA2 promoters. Accordingly, we examined the effect of FOXA2 protein isoforms on the activity of the proposed alternative FOXA2-P2 promoter. In contrast to the observed repression of the FOXA2-P1 promoter, the isoforms activated the FOXA2-P2 promoter (Fig. 2D). The opposing regulatory effects of FOXA2 on its two promoters indicate a form of counteracting auto-regulation that may serve to fine-tune overall FOXA2 transcript levels in liver cells. In silico analysis of the P1 and P2 promoter regions revealed a predicted FOXA2 binding site within the P1 promoter (albeit low-confidence), whereas no FOXA2 consensus site was detected in the P2 promoter region. This differential motif distribution suggests that FOXA2 may directly regulate transcription from P1 but not P2.

### FOXA2 potently represses transcription from viral regulatory elements

Next, we examined whether FOXA2 exerts repressive activity on viral promoters. To this end, we performed transactivation assays using five reporter constructs each of which included distinct viral regulatory elements positioned upstream of a firefly luciferase gene.

The effects were pronounced: four of the five viral elements tested were repressed by both FOXA2 isoforms (Fig. 2E). In particular, FOXA2 over-expression resulted in dramatic suppression of transcription from the cytomegalovirus (CMV) enhancer–promoter (∼85% reduction in activity) and the SV40 early promoter (∼90% reduction). To exclude the possibility that repression resulted from promoter competition or titration of limiting transcriptional machinery, FOXA2 was expressed from plasmids driven by different promoters, all of which yielded the same pattern or repression of viral elements (data not shown). These findings further substantiate our conclusion that FOXA2 functions as a transcriptional repressor and indicate that its activity is particularly robust on viral regulatory sequences, raising the possibility that FOXA2 contributes to the control of viral gene expression in liver cells.

Interestingly, both FOXA2 protein isoforms exhibited highly similar repressive effects across all viral and cellular elements tested. When considered together with the RNA-seq and transactivation analyses described above, these results indicate that both isoforms are biologically active and functionally equivalent in their regulatory capacity. Consistent with this conclusion, cycloheximide pulse-chase assays revealed little or no difference in the stability of the two FOXA2 protein isoforms (Fig. S2). Collectively, these findings suggest that the biological significance of alternative promoter usage at the FOXA2 locus is more likely to lie in transcriptional regulation of FOXA2 expression itself rather than in the generation of protein isoforms with distinct regulatory functions.

### FOXA2 binds DNA to mediate repression of viral regulatory elements

Given the pronounced repression observed at viral regulatory elements, we sought to investigate the underlying mechanism by which FOXA2 mediates transcriptional repression. Specifically, we aimed to determine whether this effect reflects an intrinsic DNA-dependent function of FOXA2—namely, repression mediated by sequence-specific binding of FOXA2 to cis-regulatory elements, either directly at the repressed regulatory element, or indirectly through transcriptional activation of downstream effectors that ultimately result in repression—or instead results from a non-specific squelching effect caused by protein over-expression (Fig. 3A). Squelching can occur when high levels of a transcription factor sequester limiting co-regulators of the transcriptional machinery, thereby diminishing their availability and indirectly suppressing transcription [31,32]. Accordingly, we generated a plasmid encoding a mutant version of FOXA2 in which three amino acids critical for its ability to bind DNA [1] —K219, S231, and W233— were substituted with alanine (A), a small, nonpolar, uncharged amino acid commonly used to disrupt side-chain–mediated interactions without introducing steric hindrance (Fig. 3B, left). Since amino acid substitutions can potentially affect protein stability [9], we assessed the expression levels of the mutant protein. Western blot analysis confirmed that transfection of HepG2 cells with the mutant FOXA2 construct resulted in protein expression levels comparable to those observed with the wild-type FOXA2 construct (Fig. 3B, right). Next, we compared the activity of wild-type FOXA2 and the DNA-binding–deficient (DBD-mutant) FOXA2 in transactivation assays. Across all four viral elements tested, the DBD-mutant failed to mediate repression (Fig. 3C). This result is inconsistent with a squelching mechanism and indicates that FOXA2-mediated repression of viral enhancer–promoter elements requires sequence-specific DNA binding. For cellular promoters, both the FOXA2 P1 and PUMA promoters exhibited a similar pattern to that observed for viral elements, with repression abolished by the DBD-mutant FOXA2 (Fig. S3). In contrast, repression of the TP53 promoter was preserved in the presence of the DBD-mutant, indicating that repression at this locus is independent of FOXA2 DNA binding and consistent with a squelching-like mechanism. Importantly, this effect was promoter-specific and was not observed for the majority of promoters tested. Subsequent analyses focused on regulatory elements whose repression was lost upon disruption of FOXA2 DNA binding, thereby isolating DNA-binding–dependent effects from squelching-like mechanisms.

**Figure 3.**
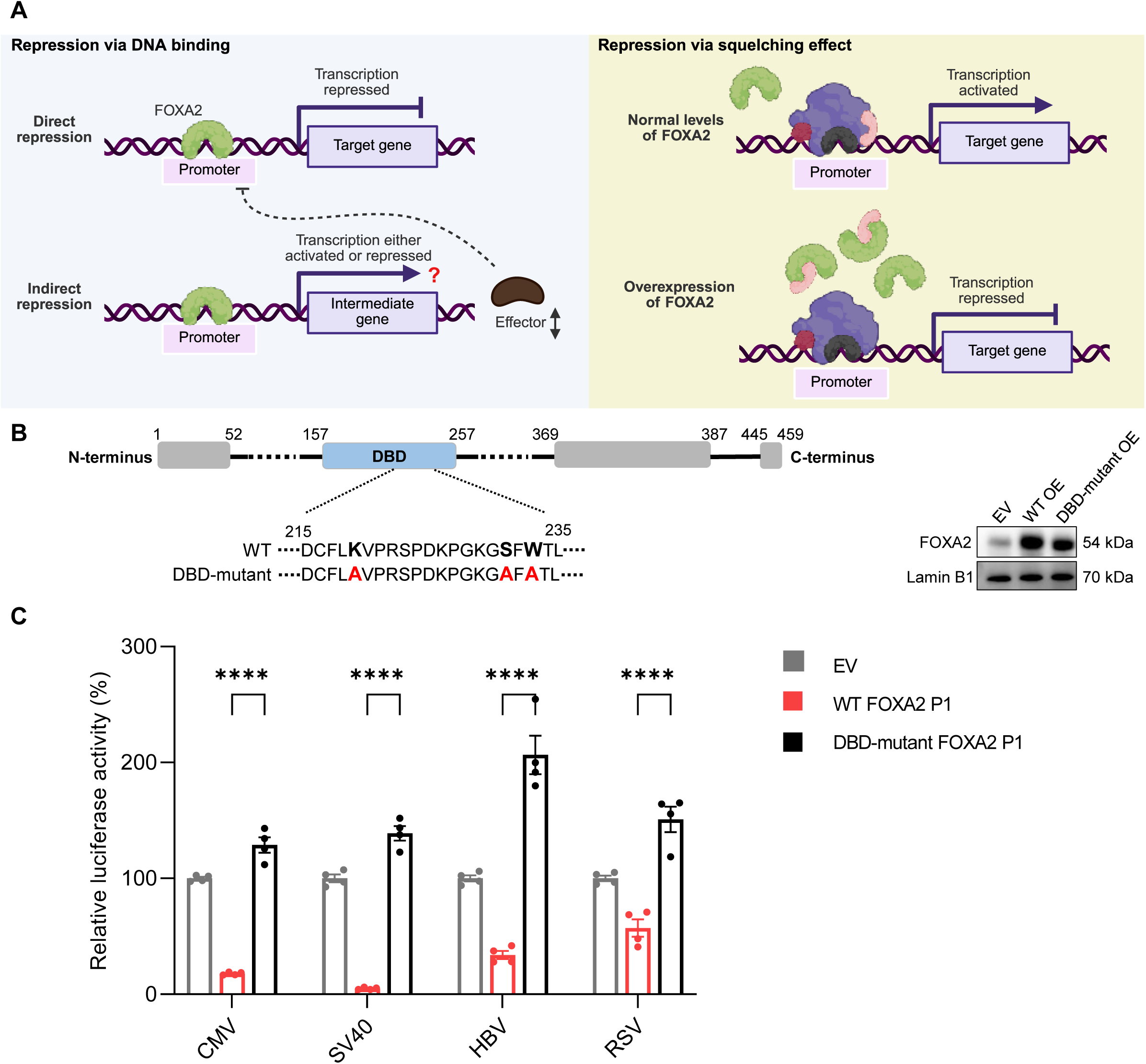
DNA binding by FOXA2 is required for repression of viral regulatory elements. **(A)** Schematic representation of alternative mechanisms by which FOXA2 could repress transcription (created with BioRender). **(B)** Schematic representation of rat FOXA2 P1 protein domains: the DNA-binding domain (DBD) is shown in blue, and the transactivation domains (TADs) are shown in grey at both termini. Numbers above the boxes indicate approximate amino acid boundaries. A zoom-in of the DBD highlights the amino acids involved in DNA recognition, with the three critical residues mutated to alanine (shown in red) in the DBD mutant. Comparison of protein stability between wild-type and mutant FOXA2 is shown in the Western blot on the right. Whole-cell protein lysates (10 µg per lane) were analyzed. The left lane represents cells transfected with empty vector (EV), the middle lane with wild-type FOXA2 P1, and the right lane with the DBD-mutant FOXA2 P1. The primary antibody recognizes the C-terminus of FOXA2 and detects both wild-type and mutant proteins. Lamin B1 was used as a loading control. Estimated molecular weights are indicated. The blot is representative of two independent experiments, each with two biological replicates. **(C)** Transactivation assays of reporter gene constructs containing viral regulatory elements by wild-type and DBD-mutant FOXA2. HepG2 cells were transiently transfected with 200 ng of plasmids containing the indicated viral elements upstream of the firefly luciferase reporter gene. In addition, 90–275 ng of either a plasmid encoding FOXA2 P1 or a plasmid encoding DBD-mutant FOXA2 P1 were co-transfected and compared with samples transfected with EV alone. The same total amount of DNA was used in all transfections (with EV used as carrier to reach 600 ng when required). Results are presented as mean ± SEM for two independent experiments with two biological replicates each. Statistical analysis was performed using ordinary two-way ANOVA with Tukey’s multiple comparisons test (****p < 0.0001).

### The SV40 early promoter is highly sensitive to FOXA2 abundance

We next focused on the viral regulatory element most strongly repressed by FOXA2, the early SV40 promoter, a well-characterized viral promoter that drives robust transcription in hepatocyte-derived cells [34]. Because the experiments described above relied on ectopic FOXA2 over-expression, which may not fully reflect physiological protein levels, we sought to validate the observed repression under conditions of reduced expression of the endogenous FOXA2 gene. To this end, we employed an RNA interference–based knockdown approach [35] to reduce endogenous FOXA2 levels and assessed SV40 promoter activity using luciferase reporter assays. HepG2 cells were transfected with siRNAs targeting both FOXA2 mRNA transcripts, resulting in ∼70% reduction in FOXA2 protein levels (Fig. 4A, upper). Importantly, this decrease was sufficient to significantly derepress the SV40 promoter, which exhibited an increase in activity of more than 1.5-fold upon FOXA2 knockdown, indicating that repression occurs also at physiological concentrations of the FOXA2 protein (Fig. 4A, lower). Other regulatory elements, viral and cellular, that were repressed by FOXA2 in a DNA-binding–dependent manner displayed only mild to moderate increases in activity following FOXA2 depletion (data not shown), underscoring the particularly strong responsiveness of the SV40 promoter to FOXA2-mediated repression. To further extend our dose–response analyses and validate the pronounced sensitivity of the SV40 promoter to FOXA2 levels, we examined promoter activity using substantially reduced amounts of FOXA2-expression plasmid, while maintaining a constant amount of SV40 reporter DNA. Strikingly, transfection with as little as 0.8% FOXA2-expressing plasmid of total transfected DNA, yielding a total FOXA2 protein level approximately three times that of endogenous FOXA2, was sufficient to induce ∼75% repression of SV40 promoter activity in HepG2 cells (Fig. 4B). Increasing FOXA2 plasmid amounts led to a modest additional decrease in promoter activity, with repression rapidly reaching a plateau at relatively low DNA concentrations. Western blot analysis confirmed a corresponding increase in FOXA2 protein levels across the tested range (Fig. 4B, upper). These findings are consistent with the siRNA-mediated knockdown experiments, and together indicate that the SV40 promoter is highly responsive to relatively small changes in FOXA2 abundance. The observation of increased SV40 promoter activity upon FOXA2 knockdown, together with the strong repression at very low FOXA2 expression levels, argue against the possibility that repression is non-physiological and only occurs under conditions of over-expression of the FOXA2 protein, and thus supports a physiologically relevant role for FOXA2 as a potent repressor of the SV40 early promoter in liver cells.

**Figure 4.**
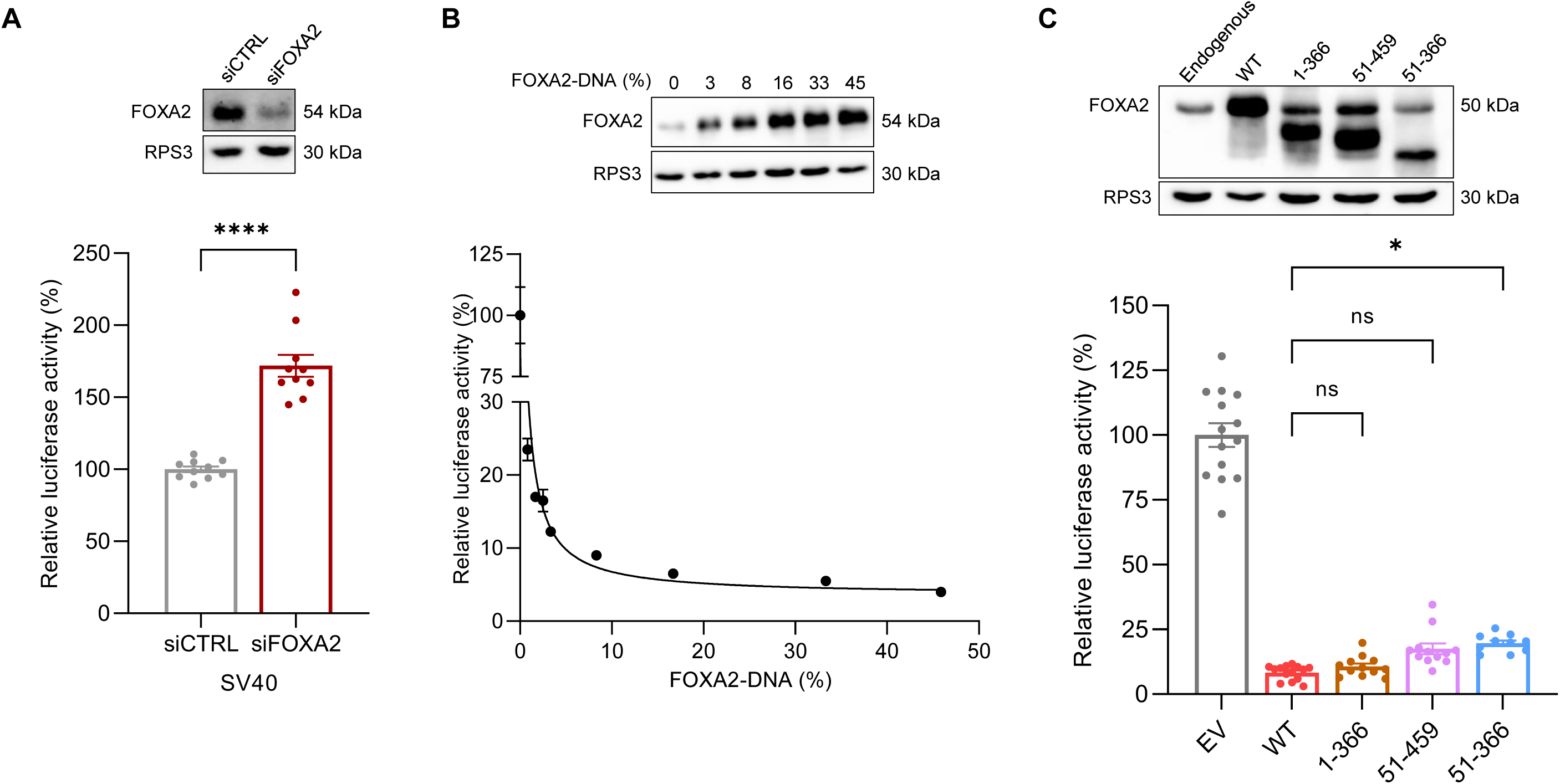
FOXA2 strongly represses the SV40 early promoter, and its DNA binding domain is sufficient to mediate this repression. **(A)** SV40 promoter activity following FOXA2 knockdown. HepG2 cells were transiently transfected with either siRNA targeting FOXA2 (siFOXA2) or a non-targeting control siRNA (siCTRL). Cells were subsequently transfected with 250 ng of an SV40 promoter–firefly luciferase reporter plasmid and 250 ng empty vector (EV). Results are presented as mean ± SEM from five independent experiments with two biological replicates each. Statistical analysis was performed using an unpaired two-tailed Student’s t test (****p < 0.0001). Western blot analysis of FOXA2 protein levels is shown above. A total of 7.5 μg protein lysate was loaded per lane. RPS3 was used as a loading control. Estimated molecular weights are indicated. The blot shown is representative of all replicates. **(B)** Dose-dependent repression of SV40 early promoter by FOXA2. HepG2 cells were transiently transfected with 275 ng of SV40 promoter–firefly luciferase reporter plasmid together with increasing amounts (5–275 ng) of a plasmid encoding FOXA2 P1. Total DNA amount was kept constant at 600 ng using EV as carrier. Data points represent the mean ± SEM from two independent experiments with two biological replicates each. Western blot analysis of selected samples confirms increasing FOXA2 protein levels corresponding to the increasing percentages of FOXA2 expression plasmid used in the transfections (7.5 μg protein per lane). **(C)** Transactivation assays of the SV40 early promoter by wild-type and truncated variants of FOXA2. HepG2 cells were transiently transfected with 200 ng of a plasmid containing the SV40 early promoter upstream of the firefly luciferase reporter gene. In addition, 90–350 ng of either noncoding EV or plasmids encoding wild-type or truncated variants of FOXA2 were co-transfected. The same total amount of DNA was used in all transfections (with EV used as a carrier to reach 600 ng when required). Results are presented as mean ± SEM from six independent experiments with two biological replicates each. Statistical analysis was performed using ordinary one-way ANOVA with Tukey’s multiple comparisons test (*p < 0.05, ns p > 0.05). Western blot analysis of 7 μg total protein per lane using an antibody recognizing an epitope proximal to the FOXA2 DNA-binding domain (DBD), confirms expression of both wild-type and truncated protein variants.

### DNA-binding domain of FOXA2 suffices for repression of the SV40 early promoter

Next, to further test whether repression of the SV40 early promoter by FOXA2 requires protein regions outwith the DNA-binding domain, we generated a series of plasmids encoding the following truncated FOXA2 variants: (i) an N-terminal fragment retaining residues 1–366, thereby lacking the C-terminal transactivation domains (TADs): (ii) a C-terminal fragment retaining residues 51–459, thereby lacking the N-terminal TAD; and (iii) a double-truncated variant lacking all known TADs, but retaining the complete DNA-binding domain. These TADs have previously been implicated in FOXA2-mediated transactivation, and their deletion is known to markedly disrupt FOXA2 interactions with transcriptional co-regulators [2,36]. Strikingly, all truncated FOXA2 variants retained strong ability to repress the SV40 early promoter (Fig. 4C, lower). Repression by the N-terminally or C-terminally truncated variants was comparable to that of the wild-type protein, whereas the DBD-only construct (51-366) exhibited slightly reduced, yet still robust, repression (75-80% decrease in promoter activity). This modest attenuation closely paralleled lower steady-state levels of the DBD-only protein detected by western blot analysis (Fig. 4C, upper). The reduced repression observed with the DBD-only construct is most simply explained by decreased protein levels, rather than a diminished intrinsic repressive capacity. The preserved repressive activity indicates that neither N- nor C-terminal transactivation domains are essential for repression of the SV40 promoter. Thus, repression of the SV40 early promoter by FOXA2 is largely mediated by its DNA-binding region further reinforcing our conclusion that sequence-specific DNA binding is the principal determinant of FOXA2-dependent repression at this viral regulatory element. Thus, DNA binding is both necessary and sufficient for repression activity.

While the SV40 early promoter was used as the primary model for mechanistic dissection of FOXA2-mediated repression, we observed that repression of additional tested regulatory elements previously shown to be repressed in a DNA-binding–dependent manner was also largely preserved with the DBD-only variant, albeit with greater variability in magnitude compared to the SV40 promoter (data not shown). Together, these data position DNA binding as the primary determinant of FOXA2-dependent repression, with additional domains fine-tuning enhancer- and promoter-specific responses.

### Identification of cis-elements underlying FOXA2-mediated repression of the SV40 promoter

To further elucidate the mechanism underlying FOXA2-mediated repression, we sought to map cis-regulatory elements within the SV40 early promoter that confer sensitivity to FOXA2. We first focused on candidate motifs previously implicated in FOXA2 or FOX-family–mediated repression. In particular, we examined TATA-like sequences within the SV40 early promoter, as FOXA2 was previously shown to bind directly to TATA box sequence to repress transcription of the ATP-binding cassette transporter A1 (ABCA1) gene [12], consistent with the similarity between the consensus TATA box motif (5’-T(A/T)T(A/T)(A/T)A(A/T)-3’;[37]) and reported FOXA2 recognition sequences (5’-T(G/A)TTT(G/A)(C/T)T-3’). In addition, another FOX-family protein, FOXP1, was reported to bind a TATA-like element within the SV40 early promoter to mediate repression [38].

Although canonical TATA box function has been reported to be largely dispensable for efficient transcription from the SV40 early promoter, two TATA-like sequences within this promoter were individually mutated, as well as in combination, to assess their potential role in FOXA2-mediated repression [39,40]. However, repression by FOXA2 was preserved in all cases, indicating that TATA box–like elements are not required for repression of the SV40 early promoter by FOXA2 (data not shown).

In parallel, to guide further analysis, we examined other promoters repressed by FOXA2 in a DNA-binding–dependent manner for shared candidate motifs. These included predicted FOX-related binding sites within the PUMA promoter and predicted FOXA binding sites within the CMV enhancer–promoter, identified using Genomatix. In addition, given a previously described mechanism in which interaction between hepatocyte nuclear factor 6 (HNF6) and the FOXA2 DNA-binding domain inhibits HNF6–DNA binding and transcriptional activity [41], we mutated the HNF6 binding site within the hepatitis B virus (HBV) enhancer–promoter and assessed repression by FOXA2. In all cases, mutation of these candidate sites failed to alleviate repression upon FOXA2 over-expression (data not shown). Collectively, these analyses demonstrate that FOXA2-mediated repression of the SV40 early promoter cannot be explained by interaction with TATA box–like sequences or known FOX-family binding motifs, pointing instead to a distinct or composite cis-regulatory architecture underlying this repression. This conclusion prompted us to next introduce large-scale deletions within the SV40 promoter, using restriction-free cloning, to localize the cis-elements responsible for FOXA2-dependent repression.

### Functional dissection of the SV40 early promoter identifies the 72-bp repeats as essential for FOXA2-mediated repression

The SV40 early promoter has been extensively characterized, with its regulatory architecture and associated transcription factor binding sites well defined [42]. It comprises two tandem 72-bp enhancer repeats at the 5′ end, followed by three 21-bp enhancer repeats—two identical and one variant—which together encompass six GC-rich motifs (5′-CCGCCC-3′) known to drive strong transcriptional activity. Approximately 20 bp downstream lies an A/T-rich region containing two TATA-like elements, followed by transcription start sites (Fig. 5A, wild-type (WT) construct). This well-defined modular organization provided a framework for systematically dissecting the promoter to identify regions required for FOXA2-mediated repression. To this end, we generated a series of deletions within the promoter (Δ1–5, Fig. 5A), each removing defined promoter elements. The relative basal activity of each deletion construct is indicated in Fig. 5A, showing that removal of any of these elements substantially reduced promoter activity. For example, removal of the 21-bp repeat region (construct Δ4) resulted in an over 10-fold decrease in basal activity, consistent with previous reports highlighting the importance of GC-rich motifs for SV40 transcriptional strength [39]. Deletion of a single 72-bp enhancer repeat (Δ1) reduced basal activity approximately fivefold, consistent with a cooperative contribution of the two repeats as described previously [43]. Each deletion construct was subsequently tested in transactivation assays to determine whether FOXA2-mediated repression was retained (Fig. 5B). Repression by FOXA2 was abolished only when both 72-bp enhancer repeats were disrupted (Δ2), whereas repression was preserved upon deletion of only one 72-bp repeat, indicating functional redundancy of this element. Repression was also retained following deletion of the 21-bp repeat region (Δ4) or the transcription start site region (Δ5), although the magnitude of repression varied depending on promoter composition. In contrast, when the promoter was reduced to the A/T-rich region containing only the TATA-like elements and downstream sequences (Δ3), FOXA2 caused an approximately sixfold activation of promoter activity. To examine whether this activation was mediated through the A/T-rich region itself, we introduced mutations within this sequence in the Δ3 construct. FOXA2-dependent activation was preserved following these mutations (data not shown), indicating that activation does not depend on specific A/T-rich sequences. Notably, basal activity of the A/T-mutated Δ3 construct was only mildly affected, consistent with previous observations that this region is dispensable for robust SV40 transcription relative to the upstream enhancer repeats (data not shown).

**Figure 5.**
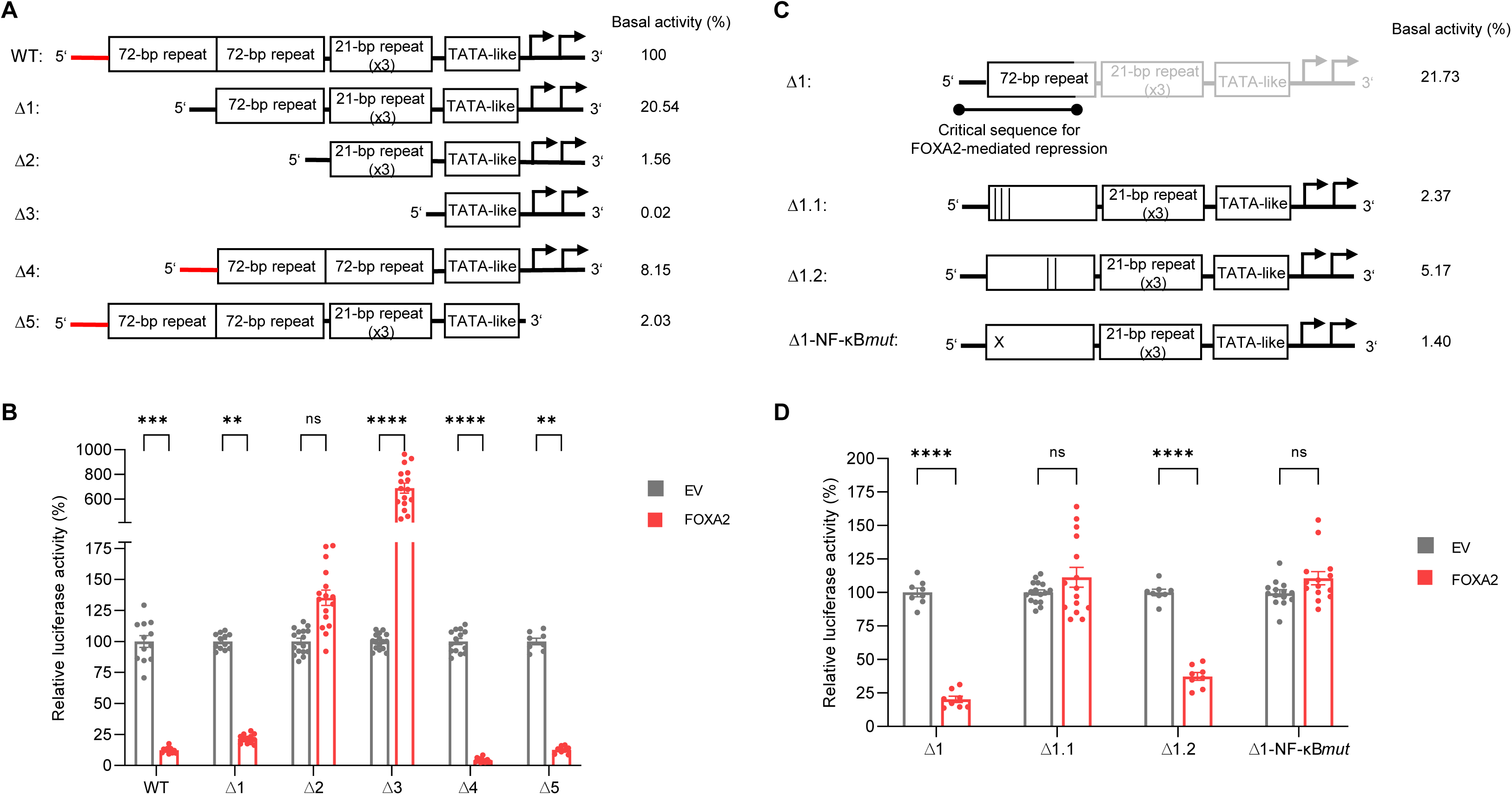
Mapping of cis-elements in the SV40 early promoter required for FOXA2-mediated repression. **(A)** Schematic representation of the SV40 early promoter, its major regulatory elements, and the deletion constructs generated. Arrows indicate multiple transcription start sites. Changes in Basal promoter activity (as a percentage of intact promoter) was measured by reporter gene assay and is indicated next to each construct. **(B)** Transactivation assays of the wild-type and deletion constructs of the SV40 early promoter by FOXA2. HepG2 cells were transiently transfected with 100–200 ng of plasmids containing either the wild-type or deletion constructs of the SV40 early promoter upstream of the firefly luciferase reporter gene. In addition, 60–90 ng of a plasmid encoding FOXA2 was co-transfected and compared with samples transfected with noncoding empty vector (EV) alone. The same total amount of DNA was used in all transfections (with EV used as a carrier to reach 600 ng when required). Results are presented as mean ± SEM from six independent experiments with two biological replicates each. Statistical analysis was performed using ordinary two-way ANOVA with Sidak’s multiple comparisons test. **(C)** Schematic representation of the Δ1 SV40 early promoter construct with site-directed mutations in the cis-element mediating repression by FOXA2. Internal deletions are indicated by vertical lines, while substitutions within the NF-κB binding site (predicted by Genomatix) are shown as “x”. Intact SV40-firefly luciferase plasmid reference and primer sequences used to generate the mutated constructs are provided in Tables S2 and S3. Basal promoter activity (as a percentage of intact promoter) is indicated next to each construct. **(D)** Same experimental setup as (B) using SV40 promoter variants shown in panel (C). (****p < 0.0001, ***p < 0.001, **p < 0.01, ns p > 0.05).

Collectively, these results identify the 72-bp enhancer repeats as the critical cis-regulatory elements required for FOXA2-mediated repression of the SV40 early promoter.

### FOXA2-mediated repression of the SV40 promoter requires an NF-**κ**B binding site within the 72-bp enhancer

These findings prompted us to further dissect the 72-bp enhancer repeats to pinpoint the cis-element required for FOXA2-mediated repression. Because the two 72-bp repeats are identical, we used the Δ1 construct, which retains a single 72-bp repeat, as a simplified background for mutational analysis, thereby avoiding the need to introduce paired mutations into the full promoter. Within this context, we generated two internal deletions targeting sequences implicated in transcription factor binding, as predicated by Genomatix and supported by previous studies of SV40 enhancer function [42] (Fig. 5C). In construct Δ1.1, a 21-bp segment at the 5′ portion of the 72-bp repeat was removed, encompassing a region predicted to bind multiple transcription factors, including NF-κB. This deletion resulted in >10-fold reduction in basal promoter activity, compared to that observed for the Δ1 construct. In contrast, construct Δ1.2 bearing a smaller 12-bp deletion that removes the Oct1 binding site retained a higher basal promoter activity. Strikingly, repression by FOXA2 was abolished in the Δ1.1 mutant, whereas it was preserved in Δ1.2 (Fig. 5D), thereby further narrowing the critical sequence required for FOXA2-mediated repression. The only well-defined consensus element within the Δ1.1 deletion corresponds to the NF-κB binding site. To directly test its involvement, we introduced a targeted base-substitution mutation disrupting the NF-κB core motif (5′-**GGGA**CTTTCC-3′ mutated to 5′-**CTTC**CTTTCC-3′; construct Δ1-NF-κB*mut*). This mutation phenocopied the Δ1.1 deletion, resulting in both a comparable reduction in basal promoter activity, and a complete loss of FOXA2-mediated repression. According to Genomatix predictions, this mutation is not expected to disrupt binding of other transcription factors within this region, with the exception of a putative hepato-nuclear factor alpha (HNF4α) binding site not previously described in the literature. The predicted HNF4α and NF-κB binding motifs largely overlap, precluding the generation of mutations that selectively disrupt one factor’s binding while preserving the other.

Although FOXA2 has been reported to interact with HNF4α in transcriptional activation contexts [44], multiple lines of evidence argue against a role for HNF4α in FOXA2-mediated repression of the SV40 promoter. HNF4α expression was unchanged upon FOXA2 over-expression (Fig. S4A–C), and although HNF4α can bind the NF-κB consensus sequence in vitro, co-expression of FOXA2 did not enhance HNF4α binding or reveal competition with NF-κB (Fig. S4D). Thus, loss of repression upon mutation of this element reflects disruption of NF-κB–dependent regulation rather than involvement of HNF4α.

Together, these data identify the NF-κB binding site within the 72-bp enhancer repeat as a critical cis-element required for repression of the SV40 early promoter by FOXA2, while also contributing to its especially high basal transcriptional activity.

### FOXA2 represses NF-**κ**B–dependent transcription in a context-dependent manner

To determine whether FOXA2-mediated repression is strictly dependent on NF-κB, we examined its effect on a panel of NF-κB–driven reporter constructs in transactivation assays. These included synthetic reporters harboring multiple NF-κB binding sites—either identical to the consensus site present in the SV40 early promoter or variant NF-κB motifs—placed upstream of minimal core promoter elements. In addition, we analyzed the HIV-1 3′ long terminal repeat (LTR), which contains NF-κB binding sites that are essential for viral transcription and pathogenicity [45] (Fig. 6A). All reporters exhibited measurable basal activity within the linear range of detection, with the 4×NF-κB reporter displaying the lowest basal activity among the tested constructs. Across all constructs, FOXA2 significantly repressed transcriptional activity (Fig. 6B), with the most pronounced effect (∼95% repression) observed for the 4×NF-κB reporter containing four sites identical to the NF-κB motif present in the SV40 early promoter. These results indicate that the magnitude of repression correlates with the extent to which promoter activity depends on NF-κB and on the specific sequence context of its binding sites.

**Figure 6.**
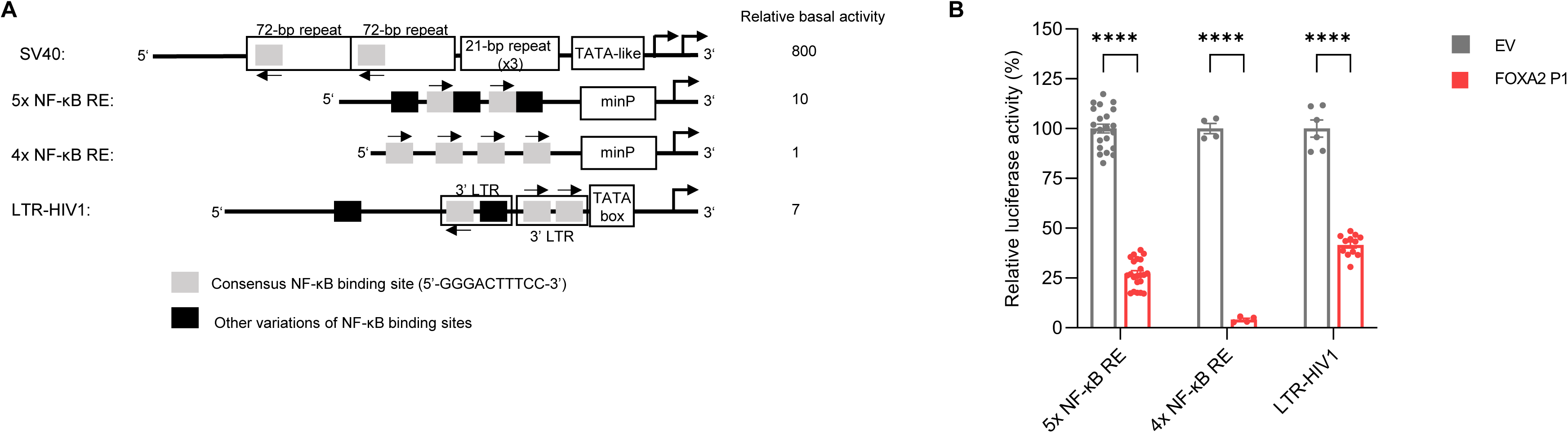
Extent of repression of NFkB-dependent transcription by FOXA2 is context-dependent. **(A)** Schematic representation of reporter constructs containing the SV40 early promoter and additional reporter constructs containing NF-κB motifs upstream of the firefly luciferase gene. Arrows above or below the boxes indicate the position of the consensus NF-κB binding sequence on the top or bottom strand, respectively, with orientation shown by the arrow’s direction (5′ → 3′). Min-P indicates minimal promoter consisting of ∼30 bp (mainly TATA box sequences), and LTR-HIV1 indicates the long terminal repeat from HIV-1. Relative basal promoter activities are shown next to each construct. **(B)** Transactivation assays of the reporter constructs shown in panel (A) by FOXA2. HepG2 cells were transiently transfected with 200 ng of the firefly luciferase reporter plasmids. In addition, 90–350 ng of a plasmid encoding FOXA2 was co-transfected and compared with cells transfected with noncoding empty vector (EV) alone. Total DNA was kept constant in all transfections (with EV used as carrier to reach 600 ng when required). Results are presented as mean ± SEM from 3–6 independent experiments with two biological replicates each. Statistical analysis was performed using ordinary two-way ANOVA with Sidak’s multiple comparisons test. (****p < 0.0001).

To further dissect this dependence, we introduced mutations into NF-κB binding sites within these regulatory elements. In the HIV-1 LTR construct, mutation of all five NF-κB sites abolished basal activity (data not shown), precluding assessment of FOXA2-mediated repression and underscoring the strict requirement for NF-κB in driving transcription from this promoter. In contrast, selective mutation of the three canonical NF-κB sites, while retaining two non-canonical motifs, reduced but did not eliminate basal activity, and under these conditions FOXA2-mediated repression was preserved (data not shown). By comparison, mutation of NF-κB sites within other previously examined regulatory elements, including the PUMA promoter and the HBV enhancer–promoter (each containing a single, non-canonical NF-κB site), as well as the CMV enhancer–promoter (containing four NF-κB sites, three of which are canonical), led to the expected reduction in basal activity, consistent with the established role of NF-κB as a transcriptional activator [46], but did not abolish repression by FOXA2 (data not shown). This may reflect differences in the relative contribution of NF-κB to basal transcription across these regulatory elements, indicating that FOXA2-mediated repression is influenced by promoter-specific regulatory architecture. Importantly, as predicted by Genomatix, the 5×NF-κB reporter does not contain any predicted HNF4α binding sites, whereas the HIV-1 LTR construct contains one. Mutation of this HNF4α binding site led to a reduction in basal promoter activity, consistent with its overlap with an NF-κB site, but did not abolish repression by FOXA2 (data not shown). Together, these observations suggest that HNF4α is unlikely to play a major role in FOXA2-mediated repression in this context. Together, these data indicate that while FOXA2 can potently repress transcription driven by NF-κB–dependent elements—particularly those highly reliant on canonical NF-κB motifs—NF-κB binding is not universally required for FOXA2-mediated repression. These findings support a model in which FOXA2 interferes with NF-κB–dependent transcription in a context-dependent manner. Accordingly, FOXA2 preferentially represses promoters that are strongly NF-κB–dependent for transcriptional output, whereas additional promoter-specific features and alternative mechanisms likely contribute to repression at other regulatory elements.

### FOXA2 represses NF-**κ**B–dependent transcription at the SV40 promoter without direct DNA binding or NF-**κ**B displacement

Next, to investigate the mechanism by which FOXA2 represses NF-κB–dependent transcription, we examined whether this effect involves changes in NF-κB expression or direct interference at the promoter level. Analysis of our RNA-seq data revealed no changes in mRNA levels of NF-κB subunits or closely associated pathway components that could account for the observed repression. Consistent with this, western blot analyses following FOXA2 over-expression did not reveal reproducible changes in protein levels of key NF-κB pathway components. Although a previous study reported reduced p-NF-κB/NF-κB and p-IKK/IKK ratios upon FOXA2 over-expression in HepG2 cells, consistent with inhibition of NF-κB signaling, these effects were observed only under oleic acid–treated conditions and were not recapitulated in our experimental system [47]. Similarly, another study reported inhibition of the NF-κB pathway by FOXA2 over-expression in lipopolysaccharide-treated hepatocytes, leading to modulation of the inflammatory response [48]. We therefore examined whether FOXA2 interferes with NF-κB binding to the SV40 early promoter. To this end, electrophoretic mobility shift assays (EMSAs) were performed using a 30-bp probe derived from the 72-bp enhancer repeat encompassing the NF-κB binding site previously identified as critical for FOXA2-mediated repression. Nuclear extracts from untreated cells yielded a weak shifted complex, approximately threefold lower in intensity than that observed following tumor necrosis factor alpha (TNFα) stimulation, likely reflecting the low basal nuclear abundance of NF-κB (Fig. 7A, lane 2). In contrast, stimulation of the canonical NF-κB pathway with TNFα, which induces nuclear translocation of p65 (RelA) and p50 (NF-κB1) [49], resulted in the appearance of a distinct DNA–protein complex consistent with NF-κB binding (Fig. 7A, lane 1). To confirm the identity of this complex, EMSAs were performed using nuclear extracts from cells over-expressing individual NF-κB subunits or defined combinations thereof. Over-expression of p50 resulted in strong DNA binding, consistent with its role as the primary DNA-binding subunit, whereas p65 alone produced only a very weak signal, in agreement with its poor intrinsic DNA-binding capacity [50,51]. Co-expression of p50 and p65 yielded a robust shifted complex comparable in mobility to that observed following TNFα stimulation, as expected, since the p50–p65 heterodimer represents the predominant transcriptionally active NF-κB complex in cells [52]. Western blot analysis (Fig. 7A, lower) confirmed over-expression of the NF-κB subunits and revealed reduced p50 protein levels when co-expressed with p65, while p65 levels remained stable or slightly increased. This is consistent with the known instability of p50, which lacks a transactivation domain and is more susceptible to degradation when not stabilized by transcriptional engagement or associated post-translational modifications, in contrast to p65 [53,54]. To further validate the specificity of NF-κB binding to this element, competition EMSAs were performed using unlabeled wild-type or NF-κB–mutant oligonucleotides. The wild-type competitor efficiently reduced binding to the labeled probe, whereas the NF-κB–mutant competitor failed to do so, confirming sequence-specific NF-κB binding. The minor reduction observed in band intensity with the mutant competitor likely reflects nonspecific DNA–protein interactions (Fig. S5).

**Figure 7.**
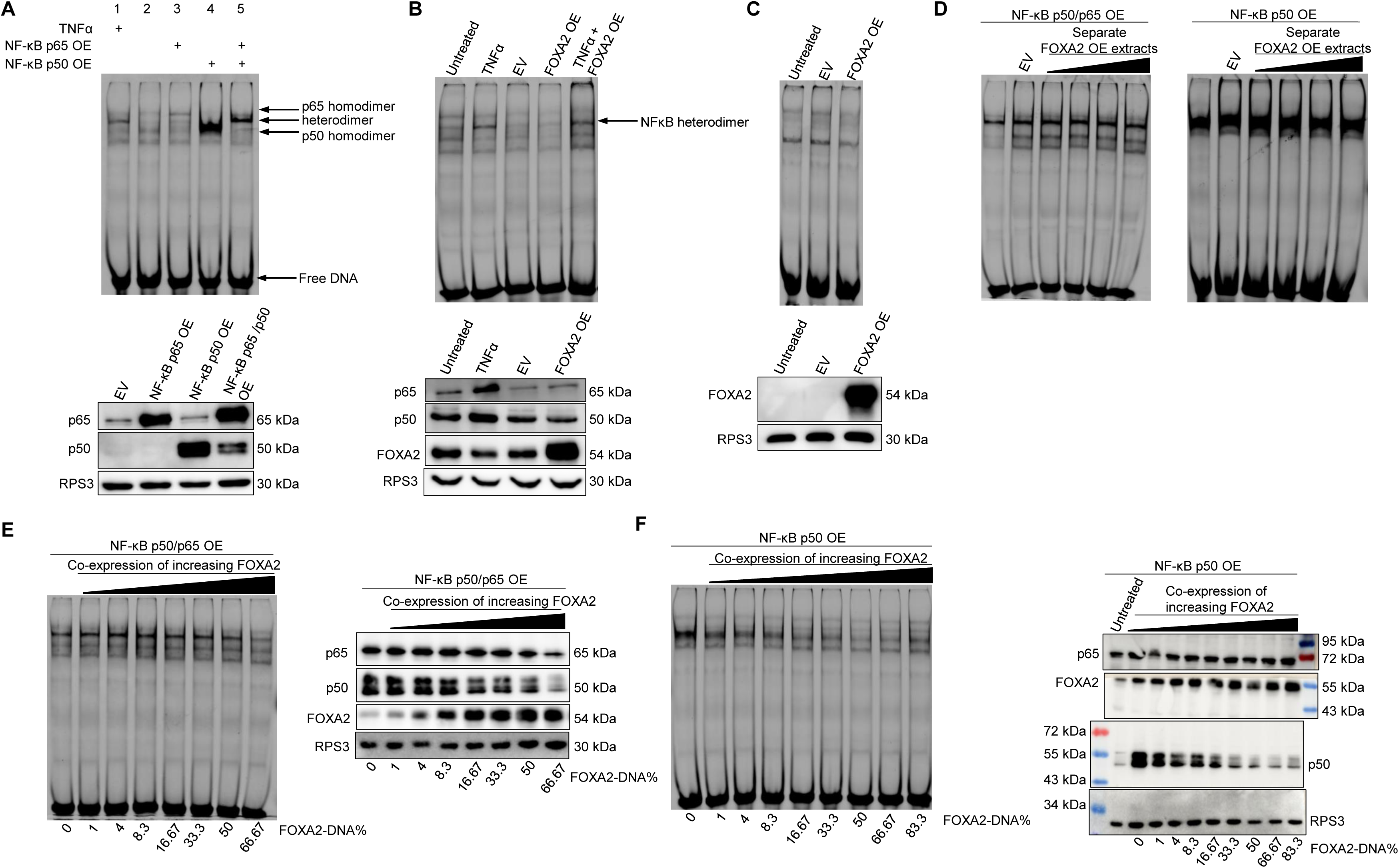
Extent of repression of the SV40 early promoter by FOXA2 is consistent with reduced NF-κB protein abundance and not with direct competition or displacement from DNA. **(A)** Upper panel: EMSA showing NF-κB–DNA complexes in nuclear extracts from HepG2 cells following TNFα treatment or over-expression of p50 and/or p65 as indicated. A fluorescently labeled double-stranded oligonucleotide corresponding to the NF-κB response element identified as critical for FOXA2-mediated repression in the SV40 early promoter was used as probe. Shifted complexes corresponding to NF-κB homo- or heterodimers are indicated by arrows. Five μg of nuclear protein were loaded per lane. Lower panel: Western blot analysis of nuclear extracts from the same samples used for EMSA. A total of 10 μg protein were loaded per lane. RPS3 was used as a loading control. Estimated molecular weights are indicated **(B)** A similar experimental setup was used, with conditions indicated above the EMSA and Western blot. For EMSA, 5 μg of nuclear protein were loaded per lane, except in the final lane, where equal amounts (5 μg each) from FOXA2 over-expression and TNFα-treated samples were mixed. For Western blot, 7.5 μg of nuclear protein were loaded per lane. **(C)** Nuclear extracts from HEK293T cells following the indicated treatments were used for EMSA (upper; 5μg of protein per lane) and Western blot (lower; 10 μg of protein per lane). **(D)** EMSA showing NF-κB–DNA complexes in nuclear extracts from HepG2 cells following over-expression of constant amounts of plasmids encoding p50/p65 or p50 alone, as indicated. In the first lane, 5 μg of nuclear extract from NF-κB–over-expressing cells was loaded alone. In subsequent lanes, these extracts were mixed with separate nuclear extracts from cells transfected with noncoding empty vector (EV) or increasing amounts of plasmid encoding FOXA2. Equal amounts of protein from each extract (5 μg + 5 μg) were combined prior to EMSA. **(E–F)** EMSA (left) showing NF-κB–DNA complexes in nuclear extracts from HepG2 cells following co-expression of increasing amounts of FOXA2 (transfected DNA percentages shown below the gel) together with constant amounts of plasmids encoding p50/p65 (panel E) or p50 alone (panel F), as indicated. In all lanes, 5 μg of nuclear extract was loaded. Western blot analysis (right) of nuclear extracts from the same samples used for EMSA. A total of 10 μg protein was loaded per lane. The Western blot in panel F shows molecular weight markers derived from two separate blots in which identical samples were loaded and processed in parallel. All results in this figure are representative of two independent experiments.

These results establish robust, sequence-specific binding of NF-κB to the SV40 72-bp enhancer repeat, providing a framework to examine whether FOXA2 interferes with this interaction directly or indirectly. We therefore first tested whether FOXA2 itself binds this sequence in vitro. As noted above, nuclear extracts from untreated cells yielded only weak bands in the EMSA, as did extracts from cells transfected with empty vector. Accordingly, we examined whether nuclear extracts from cells over-expressing FOXA2 would generate an additional shifted band corresponding to FOXA2–DNA binding. EMSA analysis revealed no detectable band attributable to FOXA2 binding (Fig. 7B, upper), consistent with bioinformatic transcription factor binding predictions, which failed to identify any high-confidence FOXA binding sites and revealed only a weak, low-confidence motif match within this region.

To test whether FOXA2 could displace NF-κB by competing on the same binding site, or by indirectly regulating other transcription factors that would, we mixed extracts from FOXA2-over-expressing cells with separate extracts from TNFα-stimulated cells. The band intensity in the mixed extract was similar to that from TNFα-treated cells alone (Fig. 7B, upper), which was confirmed by densitometric quantification, indicating no clear displacement of NF-κB from its binding site that could explain repression of the SV40 promoter in vitro. Western blot analysis of the corresponding nuclear extracts confirmed robust over-expression of FOXA2 and efficient nuclear accumulation of NF-κB subunits following TNFα stimulation (approximately 2.5-fold for p50 and 6-fold for p65; Fig. 7B, lower).

To further exclude the possibility that FOXA2 directly occupies the NF-κB binding site within the SV40 promoter, we over-expressed FOXA2 in HEK293T cells, which lack detectable endogenous FOXA2, as confirmed by western blot analysis (Fig. 7C, lower). Consistent with the results obtained in HepG2 cells, EMSA analysis revealed no detectable shifted band attributable to direct binding of FOXA2 to the DNA probe, irrespective of FOXA2 expression level or the presence of other proteins in the nuclear extract (Fig. 7C, upper).

Additionally, to test whether FOXA2 interferes with NF-κB binding to this sequence in a dose-dependent manner and to further exclude direct competition with NF-κB, nuclear extracts from cells over-expressing graded amounts of FOXA2 were mixed with extracts containing over-expressed p50/p65 heterodimers or p50 homodimers (Fig. 7D). In all conditions tested, NF-κB binding to the probe was preserved, with no reduction in band intensity correlated with increasing FOXA2 levels, as confirmed by densitometric quantification and shown in Fig. S6A. Minor band diffusion and the appearance of nonspecific complexes in some lanes are most likely due to protein overload rather than specific effects on DNA binding. Together, these results argue against a model in which FOXA2 represses the SV40 promoter by competing with or displacing NF-κB from its cognate binding site, either directly or indirectly via regulation of other transcription factors that would mediate such displacement.

### FOXA2-mediated repression of the SV40 early promoter correlates with decreased NF-**κ**B protein abundance

We next asked whether FOXA2 modulates NF-κB in vivo in a manner that attenuates its binding to the SV40 early promoter. Because untreated cells yielded only weak, barely detectable NF-κB binding in EMSA, we examined the effects of co-expressing NF-κB subunits with increasing amounts of FOXA2. Co-expression of FOXA2 with NF-κB resulted in a clear attenuation of NF-κB–DNA complexes in EMSA (Fig. 7E and F, left), indicating an intracellular inhibitory effect of FOXA2 on NF-κB. Higher levels of FOXA2 were required to reduce binding of the p50/p65 heterodimer compared with the p50 homodimer. Approximately a 50% reduction in p50 homodimer binding was observed with as little as 4% FOXA2-expressing plasmid of total transfected DNA, whereas a comparable reduction in heterodimer binding required more than 50% FOXA2-expressing plasmid (band quantification is shown in Fig. S6B). This difference is consistent with the higher stability and DNA-binding affinity of the p50/p65 heterodimer relative to the p50 homodimer [55].

Western blot analysis revealed that the reduced DNA binding observed by EMSA coincided with decreased NF-κB protein levels upon FOXA2 over-expression (Fig. 7E and F, right). The reduction in p50 levels closely mirrored the EMSA results, appearing sharper and more pronounced when p50 was expressed alone, whereas co-expression with p65 resulted in a more gradual decline in p50 abundance, consistent with a stabilizing effect of p65. In contrast, a decrease in p65 protein levels was observed only at higher FOXA2 expression levels, in agreement with the EMSA data. At first glance, these findings appear to contrast with earlier observations (Fig. 7B, lower), in which co-expression of p50 with p65 led to reduced p50 levels. We propose that these differences reflect protein-level dynamics that depend on relative expression stoichiometry. Under conditions of excess p50, unpartnered p50 may be preferentially degraded, whereas at intermediate levels, association with p65 stabilizes p50. Only at higher FOXA2 levels does a coordinated reduction of both p50 and p65 become evident. It is important to note that FOXA2 represses SV40 early promoter activity at very low expression levels (<1% of transfected DNA), as shown earlier, whereas a clear reduction in NF-κB DNA binding in EMSA was observed only at substantially higher amounts of FOXA2. This apparent discrepancy likely reflects the lower sensitivity of EMSA to detect modest changes in NF-κB activity compared with transcriptional readouts in vivo, where small alterations in NF-κB protein levels or activity can be amplified into robust effects on promoter output.

In addition, a reduction in endogenous NF-κB protein levels was not consistently observed upon FOXA2 over-expression. Although densitometric analysis of western blots in Fig. 7B indicated a partial decrease in nuclear p50 (∼50%) and p65 (∼20%) levels, this trend was not reproducible across independent experiments. Since FOXA2 transfection efficiency was incomplete (estimated at ∼60–70%), any FOXA2-dependent reduction in endogenous NF-κB protein levels would be averaged with non-transfected cells in bulk lysates, thereby substantially dampening the apparent magnitude of the effect detected by western blotting. Consequently, the actual decrease in NF-κB protein levels within FOXA2-expressing cells is likely stronger than observed but difficult to resolve reliably at the population level. For this reason, NF-κB subunits were over-expressed in subsequent experiments to enable more robust detection of FOXA2-dependent changes in protein abundance and to facilitate mechanistic interpretation.

Taken together, these results support a model in which FOXA2 represses NF-κB–dependent transcription primarily by modulating NF-κB protein levels, rather than by directly competing for or occluding NF-κB binding sites within the SV40 promoter.

### FOXA2 represses NF-**κ**B–dependent transcription by lowering NF-**κ**B protein levels rather than altering nucleocytoplasmic distribution

Since a reduction in nuclear NF-κB p50 and p65 protein levels was evident and increased with higher amounts of FOXA2, we next asked whether FOXA2 affects the nucleocytoplasmic shuttling of NF-κB. Under basal conditions, NF-κB proteins are sequestered in the cytosol by NF-κB inhibitor alpha (IκBα); upon pathway activation, IκBα is phosphorylated and degraded, allowing NF-κB to translocate to the nucleus and activate transcription of target genes, including IκBα itself, thereby establishing a negative feedback loop [56].

Consistent with our previous observations, co-expression of maximal amounts of FOXA2 with either p50 alone or with the p50/p65 heterodimer resulted in a marked reduction in nuclear NF-κB protein levels (Fig. 8A, left). Analysis of cytosolic fractions revealed a comparable reduction in NF-κB protein levels upon FOXA2 co-expression (Fig. 8A, right), closely mirroring the decrease observed in the nuclear fraction. The parallel reduction of NF-κB proteins in both compartments argues against enhanced cytosolic sequestration or nuclear export as the primary mechanism of repression. Densitometric quantification confirmed a similar extent of reduction for both p50 and p65 across nuclear and cytosolic compartments (data not shown).

**Figure 8.**
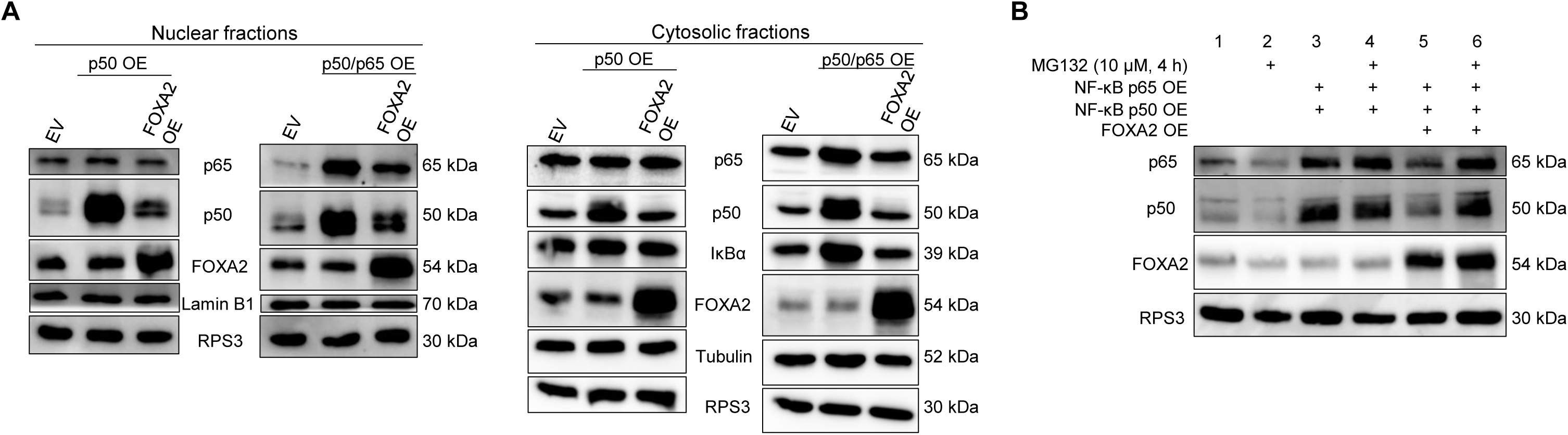
FOXA2 lowers total NF-κB protein abundance, in part by promoting proteasomal degradation. **(A)** Western blot analysis of nuclear (left) and cytoplasmic (right) fractions extracted from HepG2 cells following transfection with plasmids encoding p50/p65 or p50 alone, with or without FOXA2 co-expression, as indicated. A total of 10 μg of nuclear protein were loaded per lane, whereas an estimated equal amount of the cytoplasmic fraction was loaded (about one-third of each sample). RPS3 was used as a loading control. Efficient separation of nuclear and cytoplasmic fractions was confirmed by enrichment of Lamin B1 and tubulin, respectively. Estimated molecular weights are indicated. **(B)** To assess the contribution of proteasomal degradation to FOXA2-mediated reduction of NF-κB protein levels, Western blot analysis was performed on whole-cell extracts from HepG2 cells following over-expression of FOXA2 and NF-κB, with or without treatment with the proteasome inhibitor MG132, as indicated. Ten μg of protein were loaded per lane. Results in this figure are representative of two independent experiments.

To rule out transcriptional repression of the NF-κB expression constructs as the cause of reduced NF-κB protein levels, we examined NF-κB over-expression driven by different heterologous promoters with distinct sensitivities to FOXA2-mediated repression, including a promoter that is minimally affected by FOXA2. FOXA2 consistently reduced NF-κB protein levels to a similar extent across all constructs, even though the magnitude of transcriptional repression observed in corresponding luciferase assays was milder and varied depending on the promoter context. This disparity indicates that the pronounced decrease in NF-κB protein abundance cannot be explained by transcriptional repression of the expression vectors themselves. Instead, these results support a model in which FOXA2 acts through a post-transcriptional mechanism that directly impacts NF-κB protein levels, independently of the promoter used to drive NF-κB expression. Together, these findings indicate that FOXA2 does not primarily alter the intracellular distribution of NF-κB, but instead reduces total cellular NF-κB protein levels, thereby limiting its availability in both compartments. Consistent with this interpretation, cytosolic IκBα levels behaved as expected: over-expression of p50 alone, which lacks transcriptional activation capacity, did not induce IκBα expression, whereas over-expression of the transcriptionally active p50/p65 heterodimer increased IκBα levels, reflecting activation of the NF-κB feedback loop [52]. Notably, co-expression of FOXA2 reduced IκBα levels in this context, consistent with attenuated NF-κB transcriptional activity.

### FOXA2 promotes proteasomal degradation of NF-**κ**B proteins

Next, we investigated whether the FOXA2-mediated reduction in NF-κB protein levels involves enhanced proteasomal degradation. To test this, FOXA2 and NF-κB protein subunits were over-expressed in HepG2 cells, and NF-κB protein levels were examined in the presence or absence of the proteasome inhibitor MG132. If FOXA2 promotes proteasome-dependent degradation of NF-κB, inhibition of the proteasome would be expected to restore NF-κB protein abundance. Indeed, western blot analysis revealed that MG132 treatment substantially rescued both p50 and p65 protein levels in FOXA2-over-expressing cells, restoring them to levels comparable to those observed in the absence of FOXA2 (Fig. 8B; band quantification shown in Fig. S6C). These results indicate that FOXA2 reduces NF-κB protein levels, at least in part, through a proteasome-dependent mechanism. While these data strongly support enhanced protein degradation as a major contributor to FOXA2-mediated NF-κB suppression, they do not exclude additional mechanisms, such as effects on NF-κB protein synthesis or stability upstream of the proteasome, which were not directly addressed here.

Taken together, our results support a model in which FOXA2 represses NF-κB–dependent transcription by promoting proteasome-mediated reduction of NF-κB protein levels rather than by directly interfering with NF-κB DNA binding or nuclear localization, providing a mechanistic framework for the potent repression observed at NF-κB–sensitive promoters such as the SV40 early promoter.

## DISCUSSION

FOXA2 is a master regulator of endoderm formation and plays a central role in the maintenance of homeostasis in multiple mature organs. Consistent with this essential function, genetic ablation of FOXA2 during development is lethal [5], underscoring the need for tight and precise regulation of its expression. Genes encoding transcription factors—particularly master regulators—are frequently regulated by multiple promoters, a strategy that enables complex and context-dependent expression patterns. Alternative promoter usage provides an additional layer of regulatory control and is a major contributor to 5′ untranslated region (5′UTR) diversity, which can strongly influence mRNA stability and translational efficiency [20].

Chromatin organization varies substantially across tissues, cell types, and developmental stages, necessitating spatially and temporally controlled transcriptional regulation. In this context, alternative promoters can facilitate transcription from distinct genomic regions under specific chromatin states, thereby generating differential temporal or tissue-specific expression patterns [57,58]. A well-characterized example is GATA2, whose expression is controlled by two promoters with distinct activity profiles in hematopoietic versus non-hematopoietic tissues [59]. In other cases, alternative promoter usage generates protein isoforms with distinct functions, as exemplified by p53, in which promoter choice contributes directly to isoform diversity and functional specialization [60].

At the FOXA2 locus, we identified two transcriptionally active regions that give rise to distinct transcripts. Our data demonstrate that the non-canonical promoter is functional and indeed in the context of HepG2 cells is more active than the canonical P1 promoter. Alternative promoter usage at this locus is predicted to generate two protein isoforms that differ only at their N-termini. Direct comparison of these isoforms revealed no significant differences in protein stability, transcriptome-wide transcriptional output, or regulatory activity toward cellular and viral promoters. Consistent with these functional data, bioinformatic analysis of the six N-terminal amino acids unique to the P2 isoform (MHSASS) did not reveal similarity to known functional motifs.

While subtle isoform-specific effects cannot be excluded, particularly in vivo or under physiological conditions not captured by over-expression assays, the available evidence argues against major functional divergence between the P1- and P2-derived proteins. Instead, our results suggest that the principal biological significance of alternative promoter usage at the FOXA2 locus lies in transcriptional regulation of FOXA2 expression itself. This architecture likely enables fine-tuning of FOXA2 transcript levels in response to developmental cues, tissue-specific chromatin environments, or signaling inputs. Defining how P1 and P2 promoter usage is regulated across tissues and developmental stages will require further investigation, including analysis of chromatin state and epigenetic regulation at the FOXA2 locus.

Transcription factor abundance must be maintained within narrow limits, as deviations can have profound effects on gene regulatory networks and cell identity. Autoregulation is a common transcriptional strategy used to achieve such control, particularly for factors that occupy central positions in developmental and differentiation pathways. Autoregulatory circuits can be either positive or negative, direct or indirect, and involve simple to highly complex cis-regulatory architectures. Autoregulation can stabilize gene expression programs, sharpen temporal dynamics, and, in some contexts, lock in developmental states [29].

Our transactivation assays and RNA-seq analyses indicate that FOXA2 exhibits autoregulatory behavior in HepG2 cells. Specifically, FOXA2 activates the alternative P2 promoter while repressing transcription from the canonical P1 promoter. Negative autoregulation is a well-established mechanism for preventing excessive transcription factor accumulation and for constraining expression within an optimal range. Such circuits can influence both the amplitude and timing of gene expression, as illustrated by the Drosophila Ubx gene, which undergoes context-dependent autorepression across multiple embryonic cell types [29].

A previous report described autoactivation of the canonical FOXA2 promoter [61]. The opposing outcomes observed across studies suggest that FOXA2 autoregulation may be highly context-dependent, influenced by cellular environment, or the availability of cooperating factors. Our data argue that repression of the P1 promoter is unlikely to arise from nonspecific squelching, as repression was observed across a range of FOXA2 expression levels, and, instead, this behavior is consistent with a DNA-binding–dependent mechanism. Additionally, NF-κB binding sites were predicted by in silico analysis in the P1 promoter but not in P2, providing a potential molecular basis for FOXA2-mediated repression, including autoregulation of its own expression, as discussed in detail below.

FOXA2 has been predominantly characterized as a transcriptional activator that functions during development as a pioneer factor to access compacted chromatin, remodel nucleosomes, and facilitate gene expression [7]. Unexpectedly, our results reveal that FOXA2 can also act as a general and potent transcriptional repressor. FOXA2 over-expression led to widespread repression of endogenous genes in HepG2 cells and to strong suppression of multiple viral regulatory elements. Importantly, repression of endogenous genes may be biologically consequential, as several of these genes, including PUMA and FOXA2 itself, are implicated in the regulation of apoptosis, cell survival, and tumorigenesis. This repressive activity was supported by the effects of FOXA2 loss-of-function experiments on cellular promoters and was evident even at low FOXA2 expression levels, indicating that repression is not restricted to supraphysiological conditions. Together, these findings expand the functional repertoire of FOXA2 and challenge the prevailing view that pioneer transcription factors function exclusively as facilitators of transcriptional activation.

Viral promoters provide a particularly informative and tractable framework in which to examine transcriptional repression, as they are largely independent of higher-order chromatin context and consist of compact regulatory elements that hijack host transcription factors to optimize their life cycles in specific cellular environments [62,63]. The SV40 early promoter is a well-characterized regulatory region that relies on tandem 72-bp enhancer repeats to recruit NF-κB along with several ubiquitous transcription factors. These elements are essential for robust early gene expression, including the large T antigen (LT) critical for viral replication and cellular transformation by SV40 [64]. This clustered enhancer architecture renders the SV40 early promoter highly sensitive to changes in transcription factor activity or availability. In this context, the pronounced repression of SV40 early promoter activity by FOXA2 shown in our experimental conditions highlights both the potency and specificity of FOXA2-mediated repression.

Transcriptional repression is a critical determinant of viral persistence and latency. For example, human CMV establishes latency through chromatinization of its genome and repression of the major immediate early (MIE) promoter, a state that can be reversed by differentiation-dependent or stress-induced chromatin remodeling [65–67]. Given their ability to engage nucleosomal DNA, FOXA proteins have been proposed to facilitate viral reactivation by priming regulatory regions for transcriptional activation [68]. Our data instead support a context-dependent alternative: in differentiated liver cells, FOXA2 mediates robust repression of the SV40 early promoter, suggesting that pioneer transcription factors can also actively restrict viral gene expression depending on cellular environment and regulatory context.

Mechanistically, our data indicate that FOXA2-mediated repression of the SV40 early promoter requires direct DNA binding. Repression was abolished by mutation of the FOXA2 DNA-binding domain and was retained in truncated FOXA2 variants that preserved DNA-binding capacity but lacked TADs, identifying DNA association as a critical determinant of repression. This argues against indirect effects such as titration of transcriptional machinery or nonspecific squelching and instead supports a model in which FOXA2 directly engages regulatory DNA to repress transcription.

Directed mutagenesis of the SV40 early promoter demonstrates that NF-κB binding sites are essential both for basal promoter activity and for FOXA2-mediated repression. Disruption of these elements abolished repression, indicating that FOXA2 does not act through a nonspecific silencing mechanism but instead targets regulatory mechanisms that depend on NF-κB–driven activation. Consistent with this model, analysis of additional cellular and viral regulatory elements revealed context-dependent repression that correlated strongly with the presence and configuration of NF-κB response elements. These findings position NF-κB–dependent enhancers as a key determinant of FOXA2 repressive activity.

NF-κB is a central transcriptional regulator of inducible gene expression, controlling not only viral promoters but also a broad array of endogenous genes involved in inflammation, stress responses, proliferation, and cell survival. Dysregulation of NF-κB signaling has been directly implicated in numerous pathological conditions, including chronic inflammation and cancer [49,69]. The ability of FOXA2 to repress NF-κB–responsive regulatory elements therefore suggests a previously unappreciated mechanism by which FOXA2 may constrain excessive transcription in differentiated tissues. Mechanistically, our data do not support a model in which FOXA2 directly competes with or displaces NF-κB from DNA. Instead, the observed repression correlates with reduced NF-κB protein levels, mediated at least in part through proteasome-dependent degradation. This is supported by proteasome inhibition experiments, indicating an indirect but potent mode of regulation that operates by FOXA2-mediated regulation of the expression of an upstream intermediate gene whose expression leads to proteosome-dependent degradation of NF-κB proteins (Fig. 9). Although we cannot formally exclude the possibility that competition between FOXA2 and NF-κB for shared binding sites may occur in vivo under chromatinized conditions, or in the presence of additional cofactors, the lack of detectable competition in our in vitro assays argues that such a mechanism is unlikely to represent the primary mode of repression. How FOXA2 promotes the reduction in NF-κB protein abundance remains to be determined. Interestingly, our RNA-seq analysis revealed that FOXA2 over-expression leads to upregulation of several ubiquitin ligases that have been implicated in promoting p65 degradation, including PELI1, PDLIM4, HERC1, and HERC6 [70].

**Figure 9.**
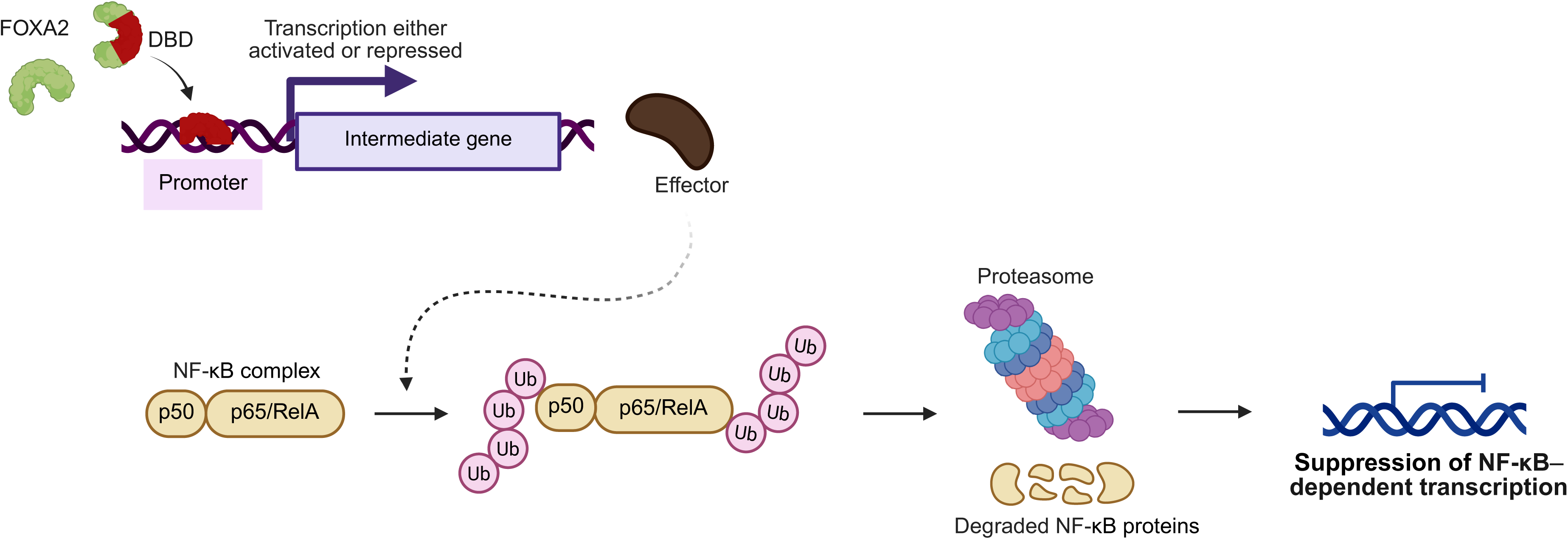
Model of FOXA2-mediated suppression of NF-κB–dependent transcription. The model (created with BioRender) illustrates that the DBD (brown) of FOXA2 (green) is sufficient to elicit a transcriptional response that promotes proteasome-mediated degradation of NF-κB, resulting in repression of NF-κB–dependent target genes.

Importantly, many viruses rely heavily on NF-κB response elements to drive early gene expression or promote the transition from latent to active infection, as exemplified by SV40 and HIV-1 [71]. These observations raise the possibility that FOXA2-mediated repression of NF-κB–driven promoters may have relevance beyond the liver, depending on tissue-specific FOXA2 expression and viral tropism. Further investigation will be required to determine the extent to which this regulatory axis operates across distinct cellular and viral contexts.

In summary, our study uncovers previously unrecognized layers of regulatory complexity at the FOXA2 locus and expands the functional repertoire of FOXA2 beyond its established role as a pioneer factor and transcriptional activator. We demonstrate that FOXA2 is expressed from alternative promoters that support distinct modes of autoregulation and that the resulting protein isoforms are functionally equivalent, suggesting that alternative promoter usage primarily serves to fine-tune FOXA2 expression rather than diversify protein function. In addition, we identify a robust repressive activity of FOXA2 that acts on both cellular and viral regulatory elements. This activity is dependent on DNA binding of FOXA2 and acts in an indirect manner on NF-κB–driven promoters including CMV and SV40 early promoter. Together, these findings highlight the context-dependent nature of FOXA2 function in differentiated cells and suggest that FOXA2 may act as a safeguard against excessive transcriptional activation. Further studies in physiological and in vivo settings will be required to determine how these regulatory mechanisms contribute to tissue-specific gene control and host–virus interactions.

## EXPERIMENTAL PROCEDURES

### Culture of human cell lines

HepG2 (human liver cancer cell line, hepatoblastoma-derived [72]) and HEK293T (human embryonic kidney cell line) were maintained at 37 °C with 5% CO_2_ in Dulbecco’s modified eagle medium (DMEM, 4.5 g/L glucose; Gibco, 41965) supplemented with 10% fetal bovine serum (FBS; Biological Industries, 04-007) and 1% Penicillin-Streptomycin solution (Corning, 30-002-Cl). For all experiments, culture plates were pre-coated with poly-L-lysine (Sigma, P4707) diluted 1:100 in phosphate-buffered saline (PBS; Sartorius, 02-023-1A) for 10 minutes at room temperature, followed by cell seeding 24-48 h prior to further treatments.

### Transient DNA transfections

To overexpress FOXA2 isoforms prior to conducting the cycloheximide-chase assay, transfections were carried out using jetPEI (Polyplus, 101000053) according to the manufacturer’s instructions. Briefly, ∼5 × 10^5^ cells were seeded per well in 6-well plates and transfected with 3 µg DNA and 6 µL JetPEI, prepared in a final of 200 µL of 150 mM NaCl solution per well. The same transfection conditions were employed for experiments with eGFP (constituting 3% of total DNA) prior to flow cytometry, as well as for experiments followed by Western blotting or electrophoretic mobility shift assays (EMSA). All transfection experiments aimed at evaluating promoter activity and FOXA2 transactivation were carried out in 24-well plates, in which 1-2 × 10^5^ cells were seeded per well and transfected with 500-600 ng total DNA, including 50 ng of an internal control plasmid in all conditions. DNA was complexed with 4 µL jetPEI in 100 µL of 150 mM NaCl solution per well. The amounts of expression and reporter plasmids used are specified in the figure legends.

### siRNA transfection

HepG2 cells were plated at ∼1.5 × 10^5^ cells per well in 24-well plates. One to two days later, cells were co-transfected with siRNA and empty plasmid DNA vector using either Lipofectamine 2000 (Thermo Fisher Scientific, 11668027) or jetPRIME (Polyplus, 101000015) according to the manufacturers’ instructions. Comparable knockdown efficiencies were obtained using Lipofectamine 2000 and jetPRIME; therefore, data generated using both reagents were combined for analysis. siRNA was used at a final concentration of 100 nM. For Lipofectamine-mediated transfections, 1 µg DNA was mixed per 30 pmol siRNA, using 1 µL Lipofectamine 2000 per 20 pmol siRNA plus an additional 2 µL per 1 µg DNA, in a total volume of 100 µL Opti-MEM (Gibco, 31985) per well. For jetPRIME-mediated transfections, identical amounts of siRNA and DNA were complexed with 4 µL jetPRIME in 100 µL jetPRIME buffer per well. Empty vector DNA was included as carrier to optimize transfection efficiency. Cells were subsequently subjected to a second round of transfection 24 h later to assess the impact on targeted promoters. siRNAs and siRNA buffer were purchased from Dharmacon: ON-TARGETplus SMART pool FOXA2 (L-010089-00-0005), ON-TARGETplus siControl non-targeting pool (D-001810-10-05), 5X siRNA buffer (B-002000-UB-100).

### Reporter gene assays

Cells in 24-well plates were lysed 24 h after plasmid transfection in 70 µL of Reporter Lysis Buffer (Promega, E397A) per well, supplemented with protease and phosphatase inhibitor cocktails, according to manufacturer’s instructions. For Renilla luciferase substrate, coelenterazine (CTZ; GoldBio, C-320) was dissolved in methanol to a stock concentration of 2.5 mg/mL, and the pH was adjusted with 0.1 N HCl. It was further diluted 1:1000 in phosphate buffer containing 80 mM K_2_HPO_4_ and 20 mM KH_2_PO_4_. Luminescence was measured in white 96-well plates (Thermo Fisher Scientific, 136101) using a Modulus microplate luminometer (Turner Biosystems), by combining 2.5-5 µL of cell lysate with 50 µL of CTZ solution. For firefly luciferase assay, 2.5-5 µL of cell lysates was combined with 25 µL of luciferin reagent (0.5 mM ATP, 33.3 mM DTT, 0.2 µg/µL co-enzyme A, 0.5 mM D-luciferin in reaction buffer [52% (w/v) (MgCO_3_)_4_·Mg(OH)_2_·5H_2_O, 20 mM tricine, 5.34 mM MgSO_4_·7H_2_O, pH 7.8]). CTZ, firefly luciferase reagents and buffer were kindly provided by the laboratory of Rivka Dikstein from the Department of Biomolecular Sciences, Weizmann Institute of Science (WIS). Renilla luciferase was expressed in all experiments under either A20 or HSV-TK promoters, and Firefly luciferase activity was normalized to Renilla luciferase activity.

### Chemical and cytokine treatments

Approximately 48 h post-transfection with FOXA2 isoforms, HepG2 cells were treated with Cycloheximide (CHX; Sigma, C7698) at a final concentration of 100 µg/mL in complete culture medium and harvested at the indicated time points. To ensure sustained inhibition of protein translation, CHX was added to the culture medium every 2 h. Cells were washed with cold PBS containing CHX and scraped on ice to maintain translational inhibition during collection. Where indicated, cells were treated with TNFα (Sigma, SRP3177) at 20 ng/mL for 30 min or with the proteasome inhibitor MG132 (Sigma, 474790) at 10 µM for 4 h. Following treatment, proteins were extracted and analyzed by SDS-PAGE followed by Western blotting or used for EMSA, as indicated.

### Protein extraction

Cells were scraped on ice, washed once with cold PBS and lysed as following: whole cell extracts were prepared by lysis in Radioimmunoprecipitation Assay Buffer (RIPA; Sigma, 20-188) supplemented with protease inhibitor (1:1000; ApexBio, K1007) and phosphatase inhibitor (1:1000; Sigma, P5726) cocktails, and centrifuged at 15,000 × g for 20 min at 4 °C. Total protein concentration was determined using the Pierce BCA protein assay kit (Sigma, 71285-3). For experiments preceded by luciferase reporter assays, RIPA buffer was added to the lysates to ensure complete disruption of all cellular membranes, including nuclear membranes. For cytosolic and nuclear fractionation, cells from 6-well plates were lysed in 150 µL Buffer A (10 mM HEPES, pH 7.9, 10 mM KCl, 0.1 mM EDTA, 0.5% IGEPAL CA-630) supplemented with protease and phosphatase inhibitor cocktails and 1 mM DTT, and centrifuged at 1,000 × g for 5 min at 4 °C. The nuclear pellet was washed once with 150 µL Buffer A lacking the detergent IGEPAL CA-630. The cytosolic fraction (supernatant) was clarified by centrifugation at 15,000 × g for 10 min at 4 °C to remove debris and membranes. Nuclear proteins were extracted by resuspending the nuclear pellet in 50 µL Buffer B (20 mM HEPES, pH 7.9, 400 mM KCl, 1 mM EDTA, 20% glycerol), supplemented with protease and phosphatase inhibitor cocktails and 1 mM DTT, followed by brief vortexing and rotation at 4 °C for 1 h. Extracts were centrifuged at 14,000 × g for 10 min at 4 °C, and supernatants containing nuclear proteins were transferred to fresh tubes. Nuclear protein concentrations were determined using the Bradford method (Bio-Rad protein dye reagent).

### Western blotting (immunoblotting)

Proteins were resolved by SDS-PAGE (10% w/v acrylamide in the separating gel) and transferred by electrophoresis to nitrocellulose membranes (Pall BioTrace, 66485). Membranes were hybridized with the following primary monoclonal antibodies diluted in 1× TBST (Bio-Lab), 5% Bovine Serum Albumin (BSA; Sigma, A7030) and 0.1% sodium azide: mouse anti-FOXA2 (Santa Cruz, sc-374376), rabbit anti-FOXA2 (Cell Signaling, 8186), mouse anti-RPS3 (Proteintech, #66046-1), rabbit anti-NF-κB1 p105/p50 (Cell Signaling, 13586), mouse anti-NF-κB p65 (Santa Cruz, sc-8008), mouse anti-IκBα (Cell Signaling, 4814), mouse anti-αTubulin (Abcam, ab7291), rabbit anti-HNF4α (Cell Signaling, 3113), and polyclonal rabbit anti-Lamin B1 (Abcam, ab16048). The secondary antibodies used were goat anti-mouse or anti-rabbit (both from Jackson ImmunoResearch, 115-035-003 and 111-035-003 respectively) IgG HRP-linked (diluted 1:10,000 in 1× TBST with 5% milk). Antibody-antigen complexes were visualized on ImageQuant 800 following enhanced chemiluminescence (ECL) exposure (Azure Biosystems, AC2103). Densitometric quantification of bands was performed using Image Lab 6.1 software (Bio-Rad) and normalized to house-keeping protein, as indicated in the results section.

### Electrophoretic mobility shift assays

Electrophoretic mobility shift assays (EMSA) were performed using a synthetic 30-mer oligonucleotide corresponding to positions -251 to -222 of the early SV40 promoter containing the NF-κB binding site (5’- CAACCAGGTGTGGAAAGTCCCCAGGCTCCC-3’). The oligonucleotide was labeled at the 5′ end with Cyanine 5 and purified by HPLC (Sigma). Double-stranded probes were generated by annealing the labeled oligonucleotide with an unlabeled complementary strand in NEB buffer (50 mM Tris, pH 7.9, 100 mM NaCl, 10 mM MgCl_2_, and 1 mM DTT final concentrations) by heating to 95 °C followed by gradual cooling to room temperature in the dark. Unlabeled wild-type and mutant double-stranded oligonucleotides were prepared in the same manner and used as competitors where indicated. For binding reactions, nuclear protein extracts (amounts indicated in the figure legends) were pre-incubated for 5 min at room temperature with binding buffer (10 mM Tris, pH 7.5, 50 mM KCl, 1 mM MgCl_2_, 0.5 mM DTT, 0.5 mM EDTA, pH 8, 200 µg/mL BSA, and 4% glycerol final concentrations) and poly(dI–dC) sodium salt (25 ng/µL; Sigma, P4929). Fluorescently labeled probe was then added to a final concentration of 10 nM in a total reaction volume of 20 µL; where indicated, unlabeled competitor oligonucleotides were added concurrently with the labeled probe. The mixture was incubated for 30 min at room temperature. Complexes were resolved from free probe by electrophoresis on a 6% (w/v) native polyacrylamide gel in 0.5× TBE buffer (Bio-Lab) at 4 °C in the dark. Gels were imaged immediately using a Typhoon fluorescence scanner directly through the gel cassettes, and band intensities were quantified by densitometry using Image Lab 6.1 software (Bio-Rad).

### Flow cytometry

HepG2 cells were harvested 36-48 hours post-transfection by incubation with 0.5% trypsin-0.2% EDTA (Bio-Lab) at 37 °C for 10 minutes, followed by gentle pipetting to ensure complete dissociation into single cells. Trypsin activity was inhibited by the addition of PBS supplemented with 10% FBS. Cells were washed and resuspended in sterile PBS containing 5% FBS and filtered into FACS tubes. GFP-positive cells were sorted using a BD FACSAria II cell sorter equipped with a 100 µm nozzle at WIS Flow Cytometry Core Facility. Sorted cells were collected into PBS containing 5% FBS, pelleted by centrifugation, and processed immediately for RNA extraction to conduct RNA-seq analysis.

### RNA-seq

Total RNA was extracted from ∼1 × 10^5^ sorted cells using NucleoSpin RNA kit (Macherey-Nagel, 740955) according to manufacturer’s instructions, with two consecutive elution steps to maximize yield. RNA integrity was assessed using TapeStation at WIS Core Facility. RNA-seq libraries were prepared according to the MARS-Seq protocol [23,24]. Single-end 75 bp reads were generated on an Illumina NextSeq platform, with an output of ∼25 million reads per sample. Library preparation, sequencing, and primary data processing were performed by The Nancy & Stephen Grand Israel National Center for Personalized Medicine at WIS (G-INCPM).

### RNA-seq analysis

Poly(A)/T stretches and Illumina adapter sequences were trimmed using Cutadapt [73]; resulting reads shorter than 30 bp were discarded. Remaining reads were aligned to the human genome (GRCh38.p13) using STAR [74] in EndToEnd mode with outFilterMismatchNoverLmax set to 0.05. Reads were assigned to RefSeq-annotated 3′ UTR regions (1000 bp), deduplicated based on unique molecular identifiers (UMIs), and gene-level counts were generated using HTSeq-count [75]. UMI counts were corrected for saturation by considering the expected number of unique elements when sampling without replacement. Differentially expressed genes were identified using DESeq2 [76] with betaPrior, cooksCutoff, and independent filtering disabled. Raw P values were adjusted for multiple testing using the Benjamini-Hochberg method [77]. Volcano plots displaying log_2_fold change versus -log_10_adjusted P values were generated using the web-based tool DE to Volcano (https://gincpm.shinyapps.io/de_to_volcano/).

### Restriction-free cloning

Restriction-free (RF) cloning was used to introduce nucleotide substitutions or deletions in the FOXA2 coding sequence and the early SV40 promoter. Briefly, a PCR product harboring the desired sequence modification with ends overlapping the destination plasmid was generated and used as a megaprimer in a subsequent PCR with the destination plasmid as template [78]. The resulting products were treated with DpnI (New England Biolabs, R0176S) for 1 h at 37 °C and transformed into *E. coli* DH5α bacteria (Bio-Lab, 959758026600). Colonies obtained were screened by Sanger sequencing at the WIS Core Facility, and full-length plasmid sequences were verified by Plasmidsaurus. All PCR reactions were performed using KAPA HiFi HotStart ReadyMix (Roche, KM2605) according to manufacturer’s instructions. Information on plasmids and primers used in this study is provided in Tables S2 & S3.

### Statistical analyses

Quantitative data are presented as the mean ± standard error of the mean (SEM), unless indicated otherwise, from independent experiments as indicated in the figure legends. Statistical analyses were performed using GraphPad Prism 9. The specific tests used are indicated in the figure legends. Statistical significance is denoted by asterisks: ns p > 0.05, *p < 0.05, **p < 0.01, ***p <0.001, and ****p < 0.0001.

## Data availability

Data are provided within the manuscript or supporting information files. The RNA-seq data generated in this study have been deposited in the Gene Expression Omnibus (GEO) under accession number GSE319902.

## Supporting information

This article contains supporting information.

## Acknowledgments

We thank Libby Kosolapov, a former member of our laboratory, for her early contributions to this project, including the generation of in-house plasmids and the initial conceptual development of FOXA2-mediated transcriptional repression. We also thank The Nancy and Stephen Grand Israel National Center for Personalized Medicine (G-INCPM) at the Weizmann Institute of Science for performing library preparation and high-throughput sequencing for the RNA-seq experiments. RNA-seq sequencing was done with advice from Revital Ronen at the Crown Genomics Institute and the analysis was assisted by Amir Szitenberg at the Mantoux Bioinformatics institute (G-INCPM).

## Funding and additional information

None

## Abbreviations

The abbreviations frequently used are

FOXA2: Forkhead box A2
SV40: Simian virus 40
NF-κB: Nuclear factor kappa-light-chain-enhancer of activated B cells
EMSA: Electrophoretic mobility shift assays
DBD: DNA-binding domain
TAD: Transactivation domain
TF: Transcription factor;
PUMA: p53 upregulated modulator of apoptosis.

