## Supplementary Figures for "FOXA2 potently represses viral gene expression by targeting NF-κB–dependent transcription in liver cells"

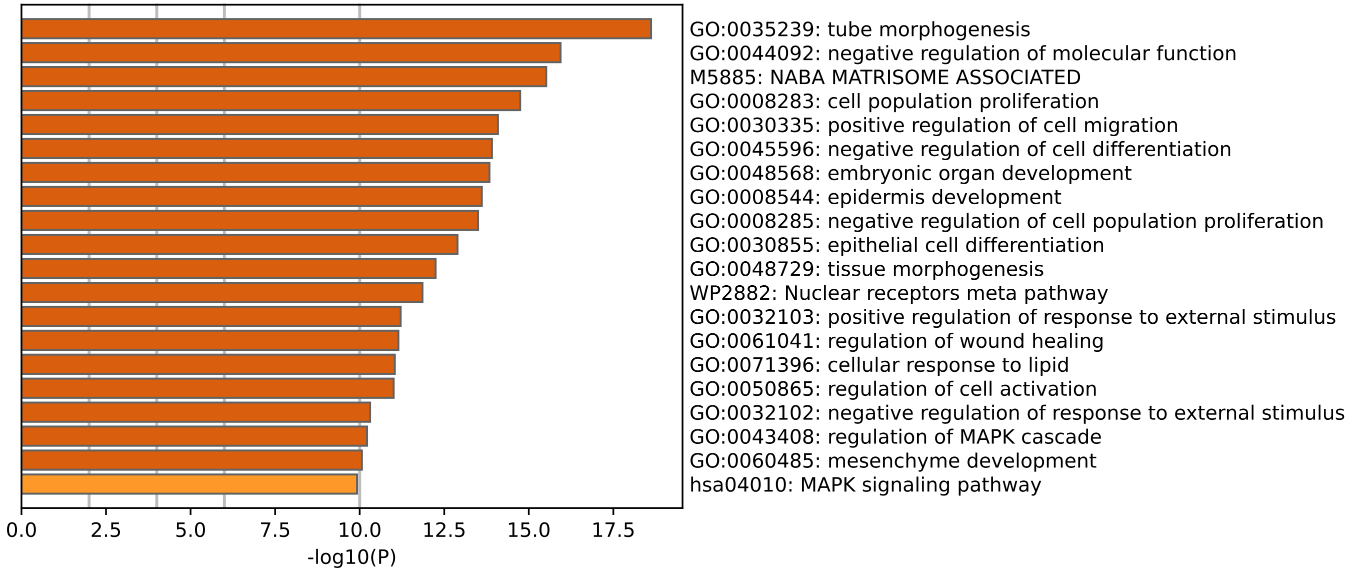


**Figure S1. Enriched pathways in HepG2 cells following FOXA2 over-expression.** Gene Ontology (GO) enrichment analysis of downregulated genes in HepG2 cells upon over-expression of FOXA2. Bar graphs represent significantly enriched biological processes, colored according to adjusted p-values, as determined by Metascape.

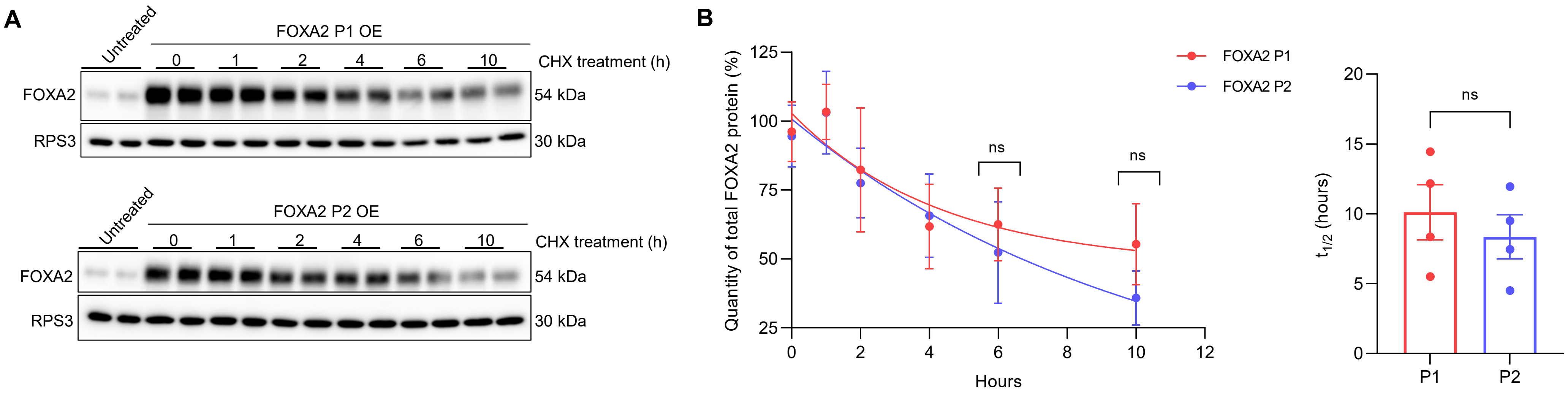


**Figure S2. Cycloheximide pulse-chase analysis of FOXA2 isoform stability in HepG2 cells.** After transient transfection of HepG2 cells with expression plasmids encoding FOXA2 protein isoforms, cells were treated with cycloheximide and harvested at the indicated time points.

**(A)** Western blot analysis of FOXA2 P1 (upper) and P2 (lower) using 7.5 µg of total protein lysate per lane, with RPS3 used as a loading control. Estimated molecular weights are indicated. “Untreated” denotes control samples not transfected with FOXA2-expressing plasmids. The blot shows biological replicates from a single experiment, which was independently repeated four times.

**(B)** Left: relative quantification of FOXA2 protein levels normalized to RPS3. Data points represent mean ± SD. Right: calculated protein half-lives based on decay curves. . Protein half-lives ($t_{1/2}$) were calculated using the equation: $t_{1/2} = \frac{\ln 2}{k_{decay}}$ , where the decay constant (k>0) was obtained by exponential decay fitting of normalized protein levels as a function of time. The estimated half-life was 10.1 h for the P1 isoform and 8.3 h for the P2 isoform. Results are presented as mean ± SEM from four independent experiments with two biological replicates each. Statistical analysis was performed using unpaired two-tailed Student’s t test (ns, $p > 0.5$).

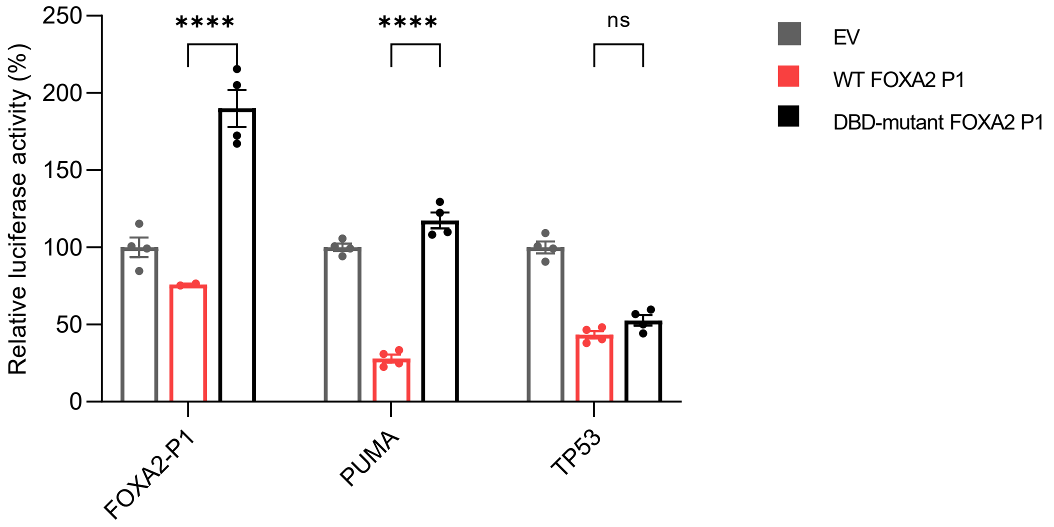


**Figure S3. DNA binding by FOXA2 is required for repression of cellular regulatory elements.**

Transactivation assays of cellular regulatory elements by wild-type and DBD-mutant FOXA2. HepG2 cells were transiently transfected with 200 ng of plasmids containing the indicated cellular promoters upstream of the firefly luciferase reporter gene. In addition, 90–275 ng of either a plasmid encoding FOXA2 P1 or a plasmid encoding DBD-mutant FOXA2 P1 were co-transfected and compared with samples transfected with noncoding empty vector (EV) alone. The same total amount of DNA was used in all transfections (with EV used as a carrier to reach 600 ng when required). Results are presented as mean ± SEM from two independent experiments with two biological replicates each. Statistical analysis was performed using ordinary two-way ANOVA with Tukey's multiple comparisons test (****$p < 0.0001$, ns $p > 0.5$).

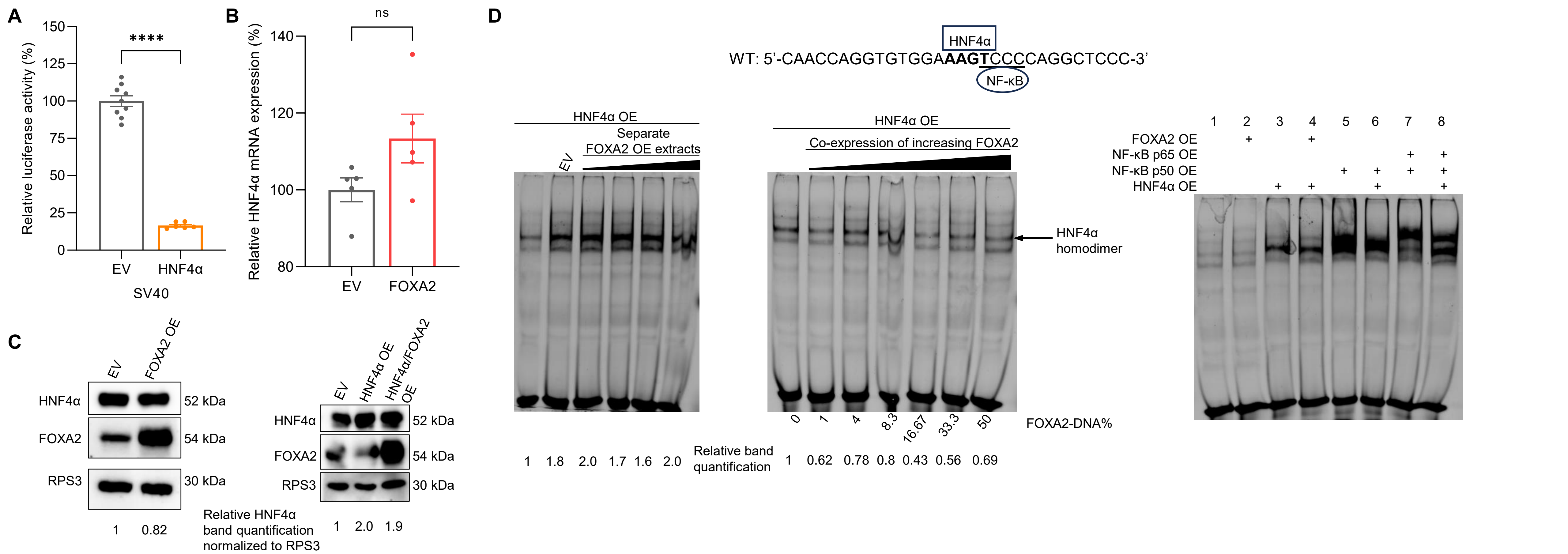


**Figure S4. FOXA2-mediated repression of the SV40 early promoter is independent of HNF4α.**

**(A)** Transactivation assays of the SV40 early promoter by HNF4α. HepG2 cells were transiently transfected with 200 ng of plasmids containing the SV40 early promoter upstream of the firefly luciferase reporter gene. In addition, 25–350 ng of a plasmid encoding HNF4α were co-transfected and compared with samples transfected with empty vector (EV) alone. The same total amount of DNA was used in all transfections (with EV used as a carrier to reach a total of 600 ng when required). Results are presented as mean ± SEM from three independent experiments with two biological replicates per experiment. Statistical analysis was performed using unpaired two-tailed Student's t test (****$p < 0.0001$).

**(B)** Relative expression levels of the endogenous HNF4α gene derived from RNA-seq analysis. Results are presented as mean ± SEM from five biological replicates. (ns $p > 0.5$)

**(C)** Western blot analysis of whole-cell (left) and nuclear (right) extracts from HepG2 cells following transfection with FOXA2 (left) or HNF4α, with or without FOXA2 co-expression (right), as indicated. A total of 10 µg of nuclear protein were loaded per lane, whereas 5 µg of whole-cell extract were loaded. RPS3 was used as a loading control. Estimated molecular weights are indicated.

**(D)** EMSA showing protein–DNA complexes in nuclear extracts from HepG2 cells following over-expression of HNF4α and NF-κB, as indicated. The transcription factor binding sites present in the oligonucleotide used for EMSA are shown in the upper panel; NF-κB and HNF4α motifs identified using the Genomatix Genome Analyzer (high-confidence binding defined as core ≥ 0.9 and matrix ≥ 0.85). **Lower left panel:** In the first lane, 5 µg of nuclear extract from HNF4α-over-expressing cells was loaded alone. In subsequent lanes, these extracts were mixed with separate nuclear extracts from cells transfected with empty vector (EV) or increasing amounts of FOXA2. Equal amounts of protein from each extract (5 µg + 5 µg) were combined prior to EMSA. **Lower middle panel:** EMSA showing HNF4α–DNA complexes in nuclear extracts from HepG2 cells following co-expression of increasing amounts of FOXA2 (transfected DNA percentages shown below the gel) together with constant amounts of HNF4α, as indicated. In all lanes, 5 µg of nuclear extract was loaded. **Lower right panel:** EMSA showing binding of NF-κB and HNF4α to DNA under the different co-expression conditions indicated above the gel. Results are representative of two independent experiments.

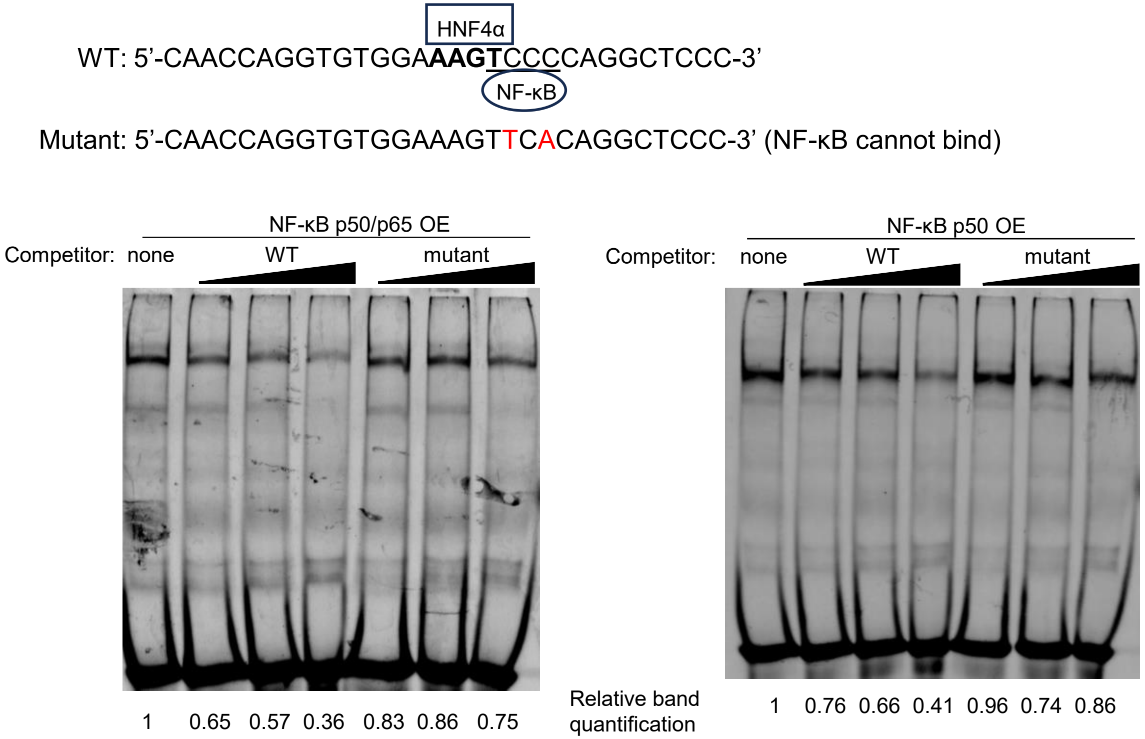


**Figure S5. Validation of NF-κB identity in EMSA by competition with wild-type and mutant oligonucleotides.**
Upper panel shows the sequences of wild-type and mutant oligonucleotides used as competitors. Lower panel shows EMSA of NF-κB–DNA complexes using nuclear extracts from cells over-expressing p50/p65 (left) or p50 alone (right). In the first lane, labeled wild-type probe was used alone. In subsequent lanes, increasing concentrations (2.5, 5, and 10 nM) of unlabeled wild-type or mutant competitor oligonucleotides were included in the binding reactions; the amount of labeled probe was identical in all samples. Five μg of nuclear extract were loaded per lane. Band quantification demonstrates specific competition of NF-κB–DNA complexes by wild-type but not mutant oligonucleotides.

**A**

| Sample | Adjusted band intensity |
|---|---|
| NF-κB p65/p50 OE | 94,677,912 |
| NF-κB p65/p50 OE + EV | 97,978,860 |
| NF-κB p65/p50 OE + extracts with increasing FOXA2 | 101,544,408 |
| | 96,713,568 |
| | 99,193,752 |
| | 100,314,774 |

| Sample | Adjusted band intensity |
|---|---|
| NF-κB p50 OE | 216,228,593 |
| NF-κB p50 OE + EV | 234,011,834 |
| NF-κB p50 OE + extracts with increasing FOXA2 | 262,552,318 |
| | 223,597,430 |
| | 217,708,834 |
| | 175,372,561 |

**B**

| FOXA2 plasmid DNA (%) | Adjusted band intensity |
|---|---|
| 0 | 48,381,760 |
| 16.67 | 40,655,494 |
| 66.67 | 14,680,512 |

| FOXA2 plasmid DNA (%) | Adjusted band intensity |
|---|---|
| 0 | 110,210,125 |
| 1 | 70,254,375 |
| 4 | 57,711,750 |
| 8.3 | 54,968,625 |
| 16.67 | 33,395,375 |
| 33.3 | 37,216,750 |
| 50 | 23,102,625 |

**C**

| | | | NF-κB p65 | | NF-κB p50 | |
|---|---|---|---|---|---|---|
| Sample | RPS3 | Normalizing factor | Band intensity | Normalized | Band intensity | Normalized |
| EV | 38,579,230 | 1.0 | 16,051,449 | 16,051,449 | 12,021,944 | 12,021,944 |
| EV + MG132 | 41,261,030 | 0.9 | 17,689,174 | 16,539,449 | 14,907,098 | 13,938,197 |
| NF-κB OE | 50,202,295 | 0.8 | 34,325,953 | 26,378,651 | 85,146,950 | 65,433,339 |
| NF-κB OE + MG132 | 52,188,994 | 0.7 | 45,635,040 | 33,734,406 | 87,558,950 | 64,725,464 |
| NF-κB & FOXA2 OE | 50,502,995 | 0.8 | 20,146,797 | 15,390,135 | 34,937,284 | 26,688,586 |
| NF-κB & FOXA2 OE + MG132 | 22,474,438 | 1.7 | 27,364,123 | 46,972,778 | 63,025,426 | 108,188,352 |

**Figure S6. Quantification of EMSA and Western blot bands.**
Tables showing densitometric quantification from a representative experiment of the gels shown in the indicated figures: **(A)** EMSA in Fig. 7D; **(B)** EMSA in Fig. 7E (upper table) and Fig. 7F (lower table); **(C)** Western blot in Fig. 8B. All densitometric quantification of bands was performed using Image Lab 6.1 software (Bio-Rad). Western blot data were normalized to the indicated loading controls. Quantifications of other experiments not shown here are available from the authors upon reasonable request.
