## Supplementary Tables for "FOXA2 potently represses viral gene expression by targeting NF-κB–dependent transcription in liver cells"

**Table S1. List of downregulated genes by FOXA2 over-expression in HepG2 cells.** A selected subset of genes that showed both the highest (negative) fold change and the lowest p-values.

| Gene | Predicted gene-associated pathways | Fold change | Adjusted p-value |
| --- | --- | --- | --- |
| ABCA12 | Lipid transport and metabolism | -28 | 9.7E-09 |
| TUBA4B | Microtubule cytoskeleton organization | -15 | 4.7E-09 |
| ALOXE3 | Lipid metabolism | -12 | 0 |
| SLC63A2 | Sphingolipid transport | -12 | 0.000000234 |
| SOCS3 | Insulin signaling | -11 | 0 |
| NEURL3 | Inhibition of hepatitis C virus assembly | -7.2 | 0 |
| NUAK2 | Glucose metabolism, Hippo pathway | -6.7 | 0 |
| ADH4 | Alcohol metabolism | -5.7 | 0 |
| EEF1A2 | Protein synthesis | -5.5 | 0 |
| GADD45B | Growth and apoptosis | -5.4 | 0 |
| AKR1B10 | Carcinogenesis | -5 | 0.00000155 |
| SLC2A3 (GLUT3) | Glucose uptake | -4 | 0 |

**Table S2. Plasmids used in this study**

| **Plasmid description** | **Source** |
| --- | --- |
| Firefly luciferase reporter driven by RSV promoter | Addgene 40343 |
| Firefly luciferase reporter driven by KSHV K8.1100 bp promoter | Addgene 131037 |
| Firefly luciferase reporter driven by human CDK4 promoter | Addgene 86656 |
| Firefly luciferase reporter driven by human TP53 promoter | Addgene 16292 |
| Firefly luciferase reporter driven by human PUMA promoter | Addgene 16591 |
| Renilla luciferase expression under HSV-TK promoter | Addgene 210571 |
| Expression of human NF-κB p65 subunit under CMV promoter | Addgene 21966 |
| Expression of human NF-κB p65 subunit under RSV promoter | Addgene 106453 |
| Expression of human NF-κB p50 subunit under CMV promoter | Addgene 21965 |
| Expression of human NF-κB p50 subunit under RSV promoter | This study |
| Firefly luciferase reporter driven by mutated CMV promoter (3 NF-κB sites mutated) | Addgene 106458 |
| Firefly luciferase reporter driven by four NF-κB response elements | Addgene 111216 |
| Expression of human HNF4α under CMV promoter | Addgene 206878 |
| Renilla luciferase expression under A20 promoter | This study |
| Expression of rat FOXA2 P1 under CMV promoter | A gift from Prof. Danielle Melloul, Hadassah-Hebrew University Medical Center, Jerusalem |
| Expression of rat FOXA2 P2 under CMV promoter | This study |
| Expression of rat FOXA2 P1 under RSV promoter | This study |
| Empty vector (EV) sharing the same backbone as the FOXA2-expressing plasmids (also called pGEM) | This study |
| Firefly luciferase reporter driven by SV40 early promoter | Bacteriology Unit, Core Facility, WIS. GenBank accession: [AY738225](http://www.ncbi.nlm.nih.gov/entrez/viewer.fcgi?db=nucleotide&val=AY738225) |
| Firefly luciferase reporter driven by CMV immediate early enhancer/promoter | Bacteriology Unit, Core Facility, WIS. GenBank accession: [EU921840](http://www.ncbi.nlm.nih.gov/entrez/viewer.fcgi?db=nucleotide&val=EU921840) |
| Firefly luciferase reporter driven by HBV inherent enhancer and promoters | Addgene 61549 |
| Firefly luciferase reporter driven by human FoxA2 P1 promoter | This study |
| Firefly luciferase reporter driven by human FoxA2 P2 promoter | This study |
| Firefly luciferase reporter lacking an upstream promoter sequence (pGL3-Basic; promoterless control) | Bacteriology Unit, Core Facility, WIS. GenBank accession: [U47295](https://www.ncbi.nlm.nih.gov/nucleotide/U47295) |
| Firefly luciferase reporter driven by five NF-κB response elements | Promega E849A |
| Firefly luciferase reporter driven by HIV-LTR. 3’-LTR fragments of HIV1 were cloned into pGL4.22 (Promega E677A) | A gift from Prof. David Wallach, Department of Biomolecular Sciences, WIS |

Mutant derivatives of the plasmids described here, generated in this study, are not listed. For plasmids without catalogue numbers, plasmid maps are available from the authors upon reasonable request.

**Table S3. Oligonucleotide primers used for RF cloning**

|  | **Mutation*** | **Primer Sequence (5’-3’)** |
| --- | --- | --- |
| **Mutations in rat FOXA2 P1 coding sequence** | K219A in DBD | F: ctctctccttcaacgactgctttctcgcggtgccccgctcgc  R: tcgcttgtgctcctgacacgga |
|  | S231A & W233A in DBD | F: cagacaagcctggcaagggcgccttcgcgaccctgcaccctgactctg |
|  | Deletion of 1-50 aa | F: ctgtgctggatatctgcagaattcagtatgggcagtggttccggcaacatgagc  R: agccgctcatgcccgccatg |
|  | Deletion of 367-459 aa | F: ctacctgcgccgccagaag  R: cagcggtttaaacttaagcttttacttcaggtgggcctctggtggcaggccag |
| **Mutations in SV40 early promoter (Fig. 5)** | ∆1 | F: ctgaaagaggaacttggttaggtacctttagtcagcaaccaggtgtggaaagtcc  R: catggtggctttaccaacagtac |
|  | ∆1.1 | F: gaacttggttaggtacctttagtcagcaacctccccagcaggcagaagtatgc |
|  | ∆1.2 | F: gtccccaggctccccagcaggcagcatgcatctcaattagtcagca |
|  | ∆1-NFκBmut | F: caaccaggtgtggaaaggaagcaggctccccagcaggcag |
|  | ∆2 | F: ctgaaagaggaacttggttaggtaccttaaccatagtcccgcccctaactccgccca |
|  | ∆3 | F: ctgaaagaggaacttggttaggtacctttaattttttttatttatgcagaggcc |
|  | ∆4 | F: gtcagcaaccatagtatggctgactaattttttttatttatgcagag |
|  | ∆5 | F: cctcggcctctgagctattccagaaaagcttggcaatccggtactgttgg  R: tatgtgcgtcggtaaaggcg |

aa=amino acid; F=forward; R=Reverse; *Where no reverse primer is listed, the same reverse primer as for the mutation in the row above was used.
